# Reporter CRISPR screens decipher *cis*- and *trans*-regulatory principles at the *Xist* locus

**DOI:** 10.1101/2024.10.08.617282

**Authors:** Till Schwämmle, Gemma Noviello, Eleni Kanata, Jonathan J. Froehlich, Melissa Bothe, Aybuge Altay, Jade Scouarnec, Vivi-Yun Feng, Martin Vingron, Edda G. Schulz

## Abstract

Developmental genes are controlled by an ensemble of *cis*-acting regulatory elements (REs), which in turn respond to multiple *trans*-acting transcription factors (TFs). Understanding how a *cis*-regulatory landscape integrates information from many dynamically expressed TFs has remained a challenge. We develop a combined CRISPR-screening approach using endogenous RNA and RE-reporters as readouts. Applied to the *Xist* locus, crucial for X-chromosome inactivation in females, this method allows us to comprehensively identify Xist-controlling TFs and map their TF-RE wiring. We find a group of transiently expressed TFs that regulate proximal REs, driving the binary activation of Xist expression. These basal activators are more highly expressed in cells with two X chromosomes, potentially driving female-specific Xist upregulation. A second set of developmental TFs is upregulated later during differentiation and targets distal REs. This regulatory axis is crucial to achieve high levels of Xist RNA, which is necessary for X-chromosome inactivation. Our findings support a model for developmental gene regulation in which factors targeting proximal REs drive binary ON-OFF decisions, while factors interacting with distal REs control the transcription output.

## Introduction

Precise and robust control of developmental genes is achieved via large and complex *cis*-regulatory landscapes^1^. In mammals, developmental genes are driven by the combined activity of up to 20 regulatory elements (REs) per gene and tissue^2–4^. Activity of each RE in turn is determined by multiple *trans*-acting signals in the form of sequence-specific DNA-binding transcription factors (TFs)^5^. Intricate TF-RE wiring thus underlies decoding of many parallel TF inputs to fine-tune gene expression. Understanding the underlying regulatory principles would necessitate simultaneously assaying a large number of TF-RE interactions, which is currently unfeasible.

While a handful of model loci have been thoroughly characterized by creating numerous mutant lines through labor-intensive genome engineering^6–8^, this approach is not readily scalable for studying a larger number of genes. Techniques to map TF-RE interactions in high-throughput instead rely on TF binding or sequence motifs^9–12^. Since they do not assess the effects on target gene or RE activity, they cannot infer regulatory relationships. A highly scalable approach to map functional TF-gene interactions are pooled CRISPR screens^13^, which however cannot identify the RE that mediates the interaction. To close this gap, we have developed a CRISPR screen variant that uses a RE reporter for phenotypic enrichment to functionally associate TFs with REs in a systematic manner.

We utilize our approach to investigate the regulation of *Xist*, an essential developmental gene with a tightly controlled and well-defined expression pattern. To trigger inactivation of one X chromosome in females, *Xist* integrates information on developmental stage and sex of the embryo. This ensures its upregulation in a female(XX)-specific and monoallelic fashion^14^. The *Xist* locus has been extensively studied for several decades, yet our understanding of how it decodes information remains limited. To identify Xist-controlling REs, we have recently performed a non-coding CRISPR screen at the onset of random X-chromosome inactivation (XCI). The screen detected a comprehensive set of 26 REs involved in initial Xist upregulation, many of which are associated with the Xist-regulating lncRNA genes *Tsix*, *Jpx*, *Ftx* and *Xert*^15^. Using chromatin state as a proxy for RE activity, we observed that distal elements exclusively respond to developmental state, whereas promoter-proximal REs are affected by X-chromosome number. This finding suggests distinct functional roles for different RE sets. The *trans*-acting factors that drive these developmental and XX-specific activation patterns remain incompletely understood.

In mice, random XCI is established during the transition from the naive to the formative pluripotent state^16–18^. Here, female cells upregulate Xist from one randomly chosen X chromosome with some cells passing through a transient biallelic state^17,19^. Notably, no developmental activators of Xist have been identified to date. Instead, the developmental dynamics of Xist are largely attributed to its repression in the naive state^20,21^. Several TFs expressed during that stage, including NANOG, OCT4 and REX1 (ZFP42) have been shown to repress Xist either directly or by promoting the transcription of Xist’s *cis*-repressor Tsix^20–24^. However, neither the deletion of *Tsix* nor the removal of the binding sites for canonical repressors induce high Xist levels prior to differentiation^25–27^. This suggests that important developmental regulators remain unidentified.

XX-specific Xist upregulation is generally believed to be driven by X-encoded Xist regulators that act in *trans*^19,28^. Two primary candidates have been identified for this role. First, the X-linked Xist activator RNF12, encoded close to the *Xist* locus, targets the repressor REX1 for degradation^23^. Importantly, removal of RNF12 during random XCI only delays, but does not prevent Xist activation, likely due to transcriptional downregulation of *Rex1* at the formative state^29,30^. Second, the lncRNA Jpx, which is encoded 10 kb upstream of *Xist*, was proposed to affect X-dosage sensing via dislodging CTCF from a promoter-proximal binding site^31,32^. However, even a combinatorial, heterozygous deletion of *Rnf12* and *Jpx* failed to abrogate Xist expression^33^. Therefore, additional X-dosage mediators are yet to be identified.

To understand how information on developmental stage and sex of the embryo is integrated at the *Xist* locus, we set out to systematically identify Xist regulators at the onset of XCI. We find a large set of previously unknown Xist activators through a pooled CRISPR interference (CRISPRi) screen, including the X-linked TF ZIC3 and the formative master regulator OTX2. Using our CRISPR screen variant that combines CRISPRi screens with a reporter assay, we systematically map TF-RE interactions across activating elements in *Xist’s cis*-regulatory landscape. We show that promoter-proximal REs are controlled by ZIC3 and a group of autosomal TFs with XX-biased expression. We propose that this regulatory axis governs initial Xist upregulation in a binary fashion to ensure inactivation of one X chromosome in females. A second group of factors, including OTX2, controls Xist expression independent of sex by interacting with distal REs. This group of activators is primarily required to achieve high transcript levels, which we show to be important for efficient X-chromosome silencing.

## Results

### Identification of Xist-controlling TFs through pooled CRISPRi screens

To comprehensively identify factors regulating Xist expression at the onset of random XCI we performed a pooled CRISPRi screen. We used differentiating mouse embryonic stem cells (mESCs) as a cell culture model to recapitulate the onset of XCI. Cells were grown in serum and 2i/LIF-containing media (2iSL), allowing efficient Xist upregulation upon differentiation by 2i/LIF(2iL)-withdrawal^15,34,35^. We further designed a CRISPRi library (TFi Lib) targeting TF genes expressed during the first 4 days of differentiation, based on the *AnimalTFDB*, a collection of mouse TFs and DNA-binding chromatin remodelers, and an RNA-seq time course experiment we had conducted previously^36,37^. In total, the TFi Lib encompassed 11,058 single guide RNAs (sgRNAs) targeting 570 TF genes and 32 non-TF genes previously implicated in Xist regulation (Extended Fig. E1a-d). Of these, we treat 17 factors, for which a loss of function has been shown to affect Xist expression, as high-confidence controls (Supplemental Table S1). We included 12 guides per target promoter, to allow the detection of subtle effects. The TFi Lib was transduced into a female mESC line carrying a split dCas9/KRAB system (TX1072 SP107), where the KRAB repressor domain is tethered to dCas9 upon addition of abscisic acid (ABA) to the media (Fig. 1a, left). To prevent X-chromosome loss, which can occur during transduction in 2iSL conditions, cells were transduced in serum/LIF (SL) media and subsequently adapted to 2iSL^15,38^. After 2 days of differentiation by 2iL-withdrawal, the cells were stained for Xist RNA using a FlowFISH assay and sorted into populations with no (Xist^Neg^), low (Xist^Low^) or high (Xist^High^) expression (Fig. 1a, Extended Fig. E1e). We sorted two different Xist-positive populations (Xist+) to be able to distinguish factors that are involved in binary control of Xist expression (“basal activators”, detected in Xist^Low^ and Xist^High^) and those required for high RNA levels (“boosting factors”, detected in Xist^High^ only). We quantified relative guide abundance in the different cell populations via sequencing of the genomically integrated sgRNA cassette. The screen displayed the expected patterns and quality control metrics (Extended Fig. E1f-i). Notably, several guides were already enriched or depleted prior to ABA induction, pointing towards leakiness of the inducible CRISPRi system (Extended Fig. E1j).

**Figure 1:**
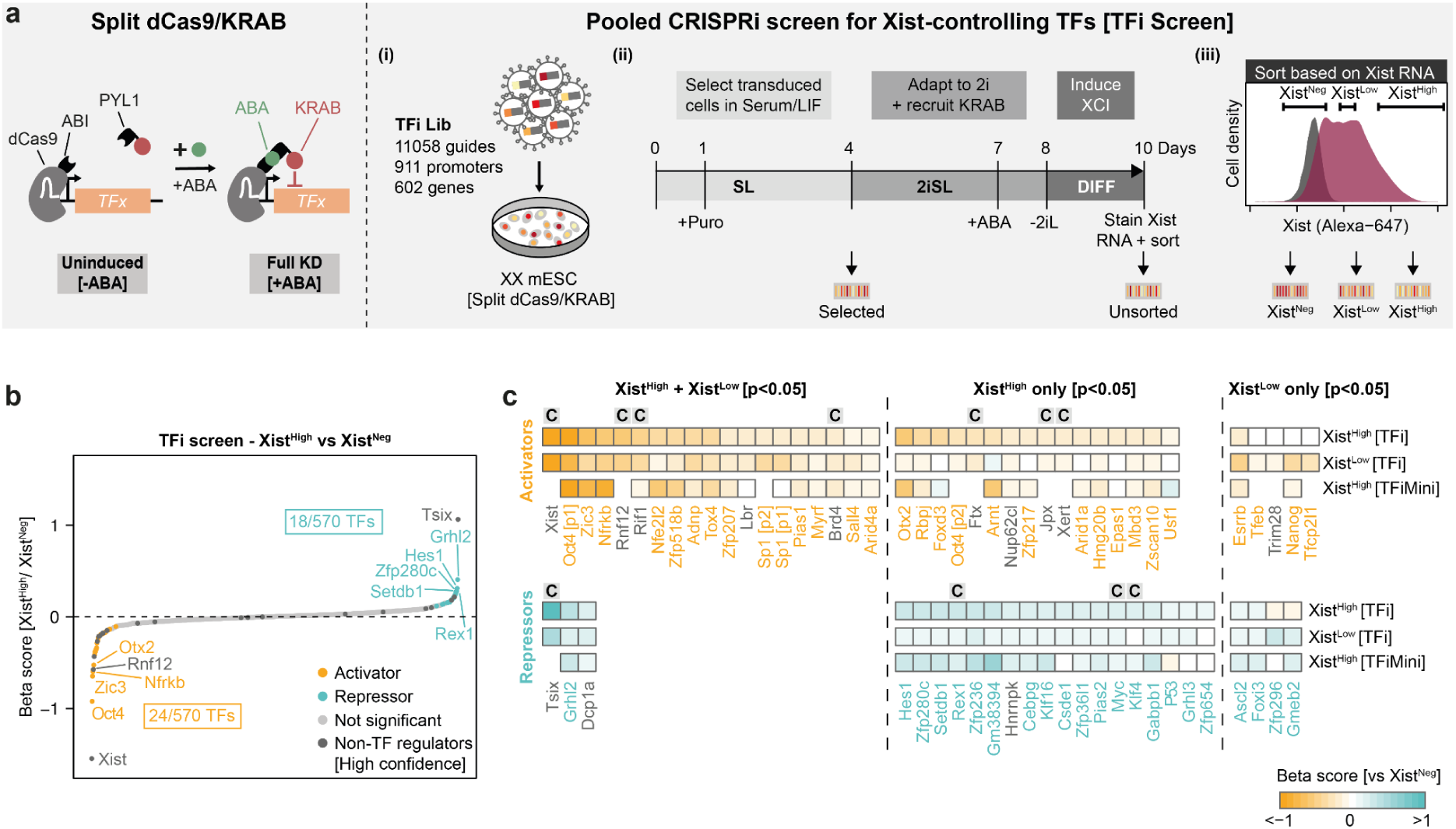
A CRISPR screen targeting expressed TFs detects novel Xist regulators. (a) Schematic of the CRISPRi system (left) and the pooled CRISPR screen (right) to identify *trans*-regulators of Xist (TFi screen). The TFi library was transduced into TX1072 XX SP107 mESCs, carrying a split dCas9-KRAB system under SL conditions (i). Following antibiotic selection and 4 days in 2iSL culture, association of the KRAB domain was induced via the addition of ABA to the medium. The cells were differentiated for 2 days via 2iL-withdrawal and stained for Xist RNA using FlowFISH (ii). Subsequently, the cells were sorted depending on Xist levels into Xist^Neg^ (bottom 15%), Xist^Low^ (bottom 15% of Xist+ cells) and Xist^High^ (top 15%) populations (iii). Relative sgRNA abundance in the different populations was assessed using next-generation sequencing (NGS). Two replicates were performed for the screen. (b) Comparison of Xist^High^ and Xist^Neg^ populations in the TFi screen. Significantly enriched or depleted target genes are colored in teal or orange respectively (*MAGeCK mle*, Wald.p≤0.05). High confidence non-TF controls are colored in dark gray (see Supplemental Table S1). (c) Heatmap depicting Xist regulators identified by comparing Xist^High^ or Xist^Low^ to the Xist^Neg^ population in the TFi screen. The bottom row compares Xist^High^ and Xist^Neg^ populations in the TFiMini screen. The plot includes all targets that were enriched or depleted in either the Xist^High^ or the Xist^Low^ population in the TFi screen (*MAGeCK mle*, Wald.p≤0.05). Orange and teal gene names indicate TFs that activate and repress Xist, respectively. High-confidence controls are marked with a “C”. Targets with empty positions in the bottom row were not included in the TFiMini screen.

To identify Xist regulators, we first compared guide abundance between the Xist^High^ and Xist^Neg^ populations (Fig. 1b, Extended Fig. E2). 11 out of 17 high-confidence regulators were enriched as expected (Wald.p≤0.05) (Supplemental Table S1). Assuringly, we observed the strongest activating and repressive effect for guides targeting the *Xist* promoter and Xist’s repressive antisense transcript *Tsix*, respectively (Fig. 1b). In addition, we also detected the known activators Rnf12, Rif1, Ftx, Xert, Jpx and the repressive pluripotency factors Rex1, Myc and Klf4^15,20,21,23,31,39^. However, our analysis did not identify other pluripotency factors, which had been reported to repress *Xist*, including Oct4, Nanog, Esrrb and Prdm14^20,21,40^. In contrast, Oct4 even scored as the strongest Xist activator in our screens. In addition, we identified 38 previously unknown Xist regulators, including 23 activators and 15 repressors (Fig. 1c, Supplemental Table S1).

For an orthogonal validation of the screen results, we performed a second smaller screen using a different CRISPRi system and a slightly adapted screen setup. We made use of our recently developed CasTuner system^41^(Extended Fig. E3a)^41^. ^41^ continuously degraded and only stabilized upon dTAG removal. We also increased the time of 2iSL adaptation from 4 to 10 days and we only sorted the Xist^High^ and Xist^Neg^ populations (Extended Fig. E3a-b). We designed a targeted sgRNA library (TFiMini Lib) encompassing 1270 guides targeting 119 putative TF regulators identified in the first screen with a permissive significance threshold (*MAGeCK mle*, Wald.p≤0.2) and 21 controls (Supplemental Table S1, Extended Fig. E3c). The TFiMini screen displayed the expected quality control metrics (Extended Fig. E3d-g) and correlated well with the TFi screen results (Peason correlation r=0.8, Extended Fig. E4a-d). Contrary to the dCas9-KRAB system, we did not observe leakiness for the inducible CasTuner system (Extended Fig. E4e-f). In the validation screen, 12 out of 24 activators and 7 out of 18 repressors initially identified in the TFi screen were confirmed (*MAGeCK mle*, Wald.p≤0.05, Extended Fig. E4c-d). Moreover, the screen identified 2 additional activators and 6 more repressors (Extended Fig. E4c-d), but the results of the two screens were highly correlated (Pearson correlation r=0.8, Extended Fig. E4b).

To classify the identified regulators according to their mode of Xist regulation, we compared guide enrichment in the Xist^High^ and Xist^Low^ populations relative to Xist^Neg^ cells in the TFi screen (Fig 1c, Extended Fig. E2c). A set of 11 TF activators were exclusively detected in the Xist^High^ population, suggesting that they primarily boost Xist levels, while 13 activators were enriched in both comparisons and might thus, in addition, control Xist in a binary fashion (Fig. 1c). The strongest activators in the first group included Oct4, the X-linked TF Zic3 and the INO80 subunit Nfrkb, while the strongest regulator in the boosting group was the epiblast regulator Otx2 (Fig. 1c)^20,42–44^.

### The canonical Xist repressor OCT4 acts as an activator during differentiation

Although previously described as a repressor, our screens identified OCT4 as a potent activator of Xist^20,22^. Notably, the repressive effect of OCT4 had been observed under different differentiation conditions or in undifferentiated cells. To disentangle these seemingly contradictory findings, we set out to characterize the role of OCT4 in Xist regulation in more detail.

We used our CasTuner cell line to repress *Oct4* in two undifferentiated (2iSL, SL) and three differentiating (-2iL, RA, EpiLC) conditions, under which Xist regulation had previously been investigated (Fig. 2a, Extended Fig. E5a-b). Xist was upregulated 13-93-fold in undifferentiated mESCs following *Oct4* knockdown (Fig. 2b-d, Extended Fig. E5c), confirming a repressive role of OCT4 under these conditions^20^. Repression of OCT4 during differentiation, by contrast, led to reduced Xist levels in all tested conditions, supporting an activating role (Fig. 2b-d, Extended Fig. E5d-e). In agreement with these results, rapid OCT4 downregulation during RA differentiation is associated with reduced Xist levels compared to 2iL-withdrawal (Extended Fig. E5a)^45^. An activating role is further supported by our previous finding that OCT4 binds several distal enhancer elements of Xist in the *Xert* region, specifically upon differentiation. In this context, OCT4 co-localizes with OTX2, which our screen identified as another strong Xist activator^15,42^. We thus propose that OCT4 acts as a direct activator of Xist during differentiation. In contrast, the repressive effect in the naive pluripotent state is potentially indirect and might be mediated by upregulation of GATA TFs upon *Oct4* depletion^40,41,46^.

**Figure 2:**
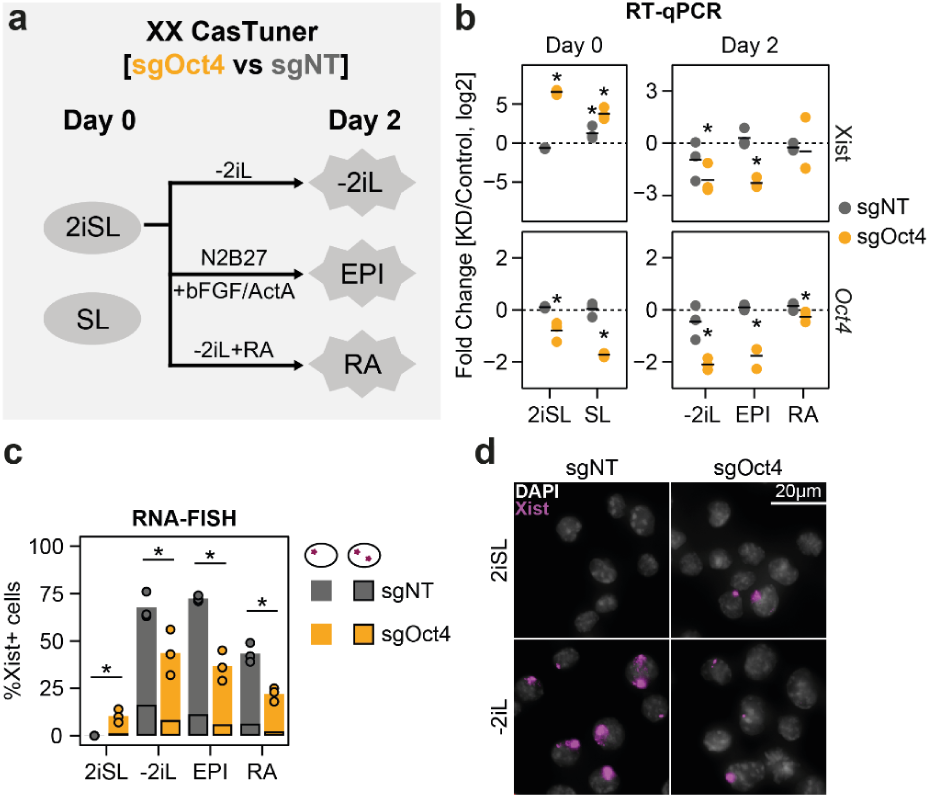
OCT4 acts as an Xist activator during differentiation. (a) Schematic outline of the experimental setup used to investigate the role of *Oct4* in Xist regulation. Knockdown of *Oct4* was induced two days before differentiation via the removal of dTAG. bFGF basic fibroblast growth factor; ActA Activin A; RA retinoic acid. (b) Effect of *Oct4* knockdown on Xist and *Oct4* expression, assessed via real-time quantitative PCR (RT-qPCR). The results are shown relative to non-targeting sgRNA (sgNT). The experiment was performed in three biological replicates. Significance between control and knockdown is marked with a black asterisk (*unpaired t-test*, p≤0.05). (c-d) Effect of *Oct4* knockdown on the percentage of Xist+ cells, assessed via RNA fluorescence *in situ* hybridization (RNA-FISH). 100 cells were counted manually per condition in c and example images for 2iSL and -2iL conditions are shown in d. The experiment was performed in 3 biological replicates and significance between the sgNT and sgOct4 samples for the total percentage of Xist+ cells was assessed using an *unpaired t-test* (p≤0.05, asterisks).

### Xist activators are transiently upregulated at the onset of XCI

Next, we aimed to systematically dissect the specific roles of a broader set of regulators in Xist regulation. Given that the *Xist* gene is upregulated specifically in females at a defined time during development^14^, we investigated how the identified Xist regulators were affected by X-chromosome number and developmental state. We performed a high-resolution RNA-seq time course experiment to assess global expression dynamics in XX and XO cells, during the first 96 hours of 2iL-withdrawal (Fig. 3a). To understand how the gene expression dynamics of our *in vitro* protocol compared to those *in vivo*, we co-embedded the data set with published scRNA-seq data covering four developmental stages from sex-mixed mouse embryos (Fig. 3b, Extended Fig. E6a)^47,48^. Our differentiation protocol captured the developmental trajectory of the embryonic tissues between embryonic days E3.5 and E6.5. Consistent with previous reports, *in vitro* differentiation of XX cells was slightly delayed in comparison to XO cells (Fig. 3b)^49,50^. Initial upregulation of Xist occurred around 24 hours after 2iL-withdrawal (Fig. 3c), corresponding to a time point between E4.5 and E5.5 *in vivo*. At this time Xist expression is also first detected by RNA-FISH according to previous work^17,19,34^. We conclude that the *in vitro* differentiation system can reproduce key aspects of developmental gene regulation around the onset of XCI.

**Figure 3:**
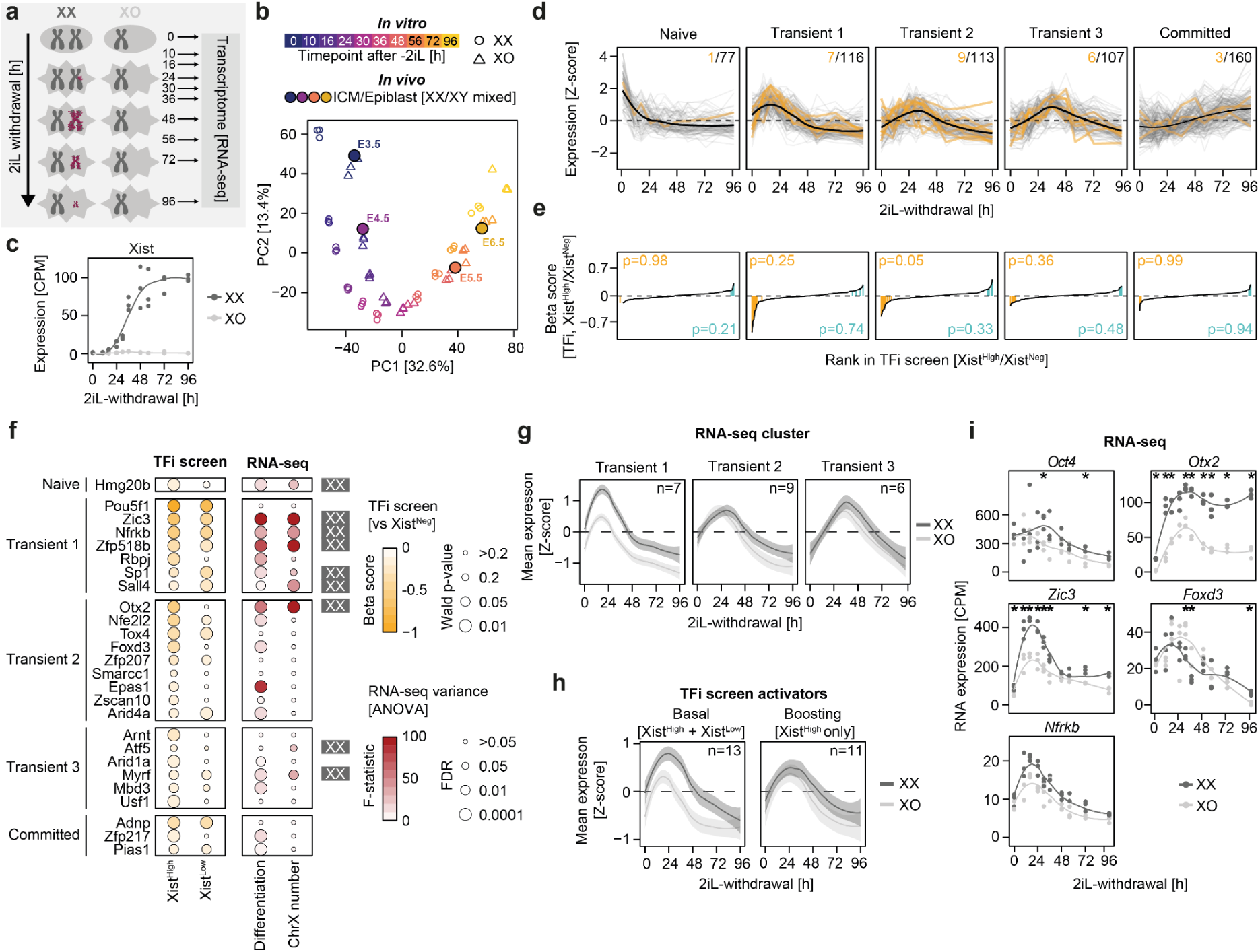
Different groups of Xist activators respond to X-chromosome number and differentiation cues. (a) Scheme depicting the experimental outline of a 2iL-withdrawal time course analyzed in b-i. Harvesting time points (in hours after 2iL-withdrawal) are indicated. The experiment was performed in three biological replicates. Xist RNA (burgundy) upregulation and spreading along the X chromosome as well as X-chromosome activity (chr. size) across the time points is indicated. (b) Joint embedding of the 2iL-withdrawal RNA-seq time course (blank shapes) with sex-mixed *in vivo* scRNA-seq data of developing mouse embryos (filled circles)^48^. Single cells from the respective time points (E3.5-E6.5) were filtered for ICM and epiblast cells and aggregated in pseudo-bulk. (c) RNA expression dynamics for Xist in XX and XO cells. (d) Clustering of z-score transformed XX expression dynamics for all TF genes included in the TFi screen (grouping based on XX and XO cells). Xist activators are shown in orange. The total numbers of genes and activators per cluster is indicated in the top right. (e) Ranked plot depicting beta scores from the TFi screen (Xist^High^/Xist^Neg^) for the TF genes included in each cluster. Activators and repressors (Wald.p≤0.05) are indicated in orange and teal, respectively. The p-values in the corner indicate enrichment of activators (orange) or repressors (teal) in the respective clusters according to a *one-sided Fisher’s exact test*. (f) Dotplot summarizing the TFi screen results and RNA-seq dynamics for all Xist activators identified in the TFi and TFi Mini screens. Statistics (Wald.p-value) and effect size (Beta score), when comparing the indicated Xist+ population to the Xist^Neg^ population (*MAGeCK mle*) in the TFi screen are shown on the left. Contribution of differentiation or X-chromosome number to the expression variance in the RNA-seq time course (*two-way ANOVA*) are shown on the right. The directionality for the X-chromosome effect is indicated next to the plot (FDR≤0.05). Expression clusters of the different activators are indicated on the left. (g) Z-score transformed RNA-seq expression dynamics of Xist activators in the transient expression clusters for XX and XO cells. The thick lines display a smoothed average across all genes in the cluster (loess method). The shaded backdrop corresponds to a 95% confidence interval. (h) Same as in (e), but grouped by the behavior of the Xist activators in the TFi screen. (i) RNA expression dynamics of selected Xist activators. Differential expression between XX and XO cells per time point is indicated by a black asterisk (*DESeq2*, Wald.FDR<0.05)^51^.

To assess the expression dynamics of the identified Xist regulators, we first grouped all TF genes targeted by the TFi Lib according to their expression patterns (naive/transient/committed) and then tested which clusters contained Xist regulators identified either in the TFi or the TFi Mini screen (Fig. 3d, Extended Fig. E6b). As expected, Xist repressors were most strongly enriched in the naive cluster (Fig. 3e, Extended Fig. E6b, p=0.21, *Fisher’s exact test*), which is downregulated when Xist is upregulated. Xist activators, on the other hand, were almost exclusively present in the three transient clusters (Fig. 3d-e, p=0.05-0.36). Notably, 7 activators, including the top-scoring TFs Oct4, Zic3 and Nfrkb, were part of the earliest transient cluster (Transient 1), which peaked before 24h and was downregulated while Xist was being upregulated. We noticed that 6 out of 7 early activators controlled basal Xist upregulation (enriched in X^High^ and Xist^Low^ in TFi screen, Fig. 3f, left). To systematically assess which factors were affected by differentiation and X-chromosome number, we performed an *ANOVA* analysis (Fig. 3f right, Extended Fig. E6c). Out of 26 identified Xist activators, 18 responded significantly to differentiation, while 9 were affected by X-chromosome number (*ANOVA*, FDR≤0.05, Fig. 3f).

Interestingly, the majority of activators in the earliest cluster (Transient 1) exhibited XX-biased expression, while the regulators in the later clusters (Transient 2/3) did not (Fig. 3f-g, Extended Fig. E6d). This pattern was also reflected in the two activator groups from the TFi screen: Basal activators were expressed in an XX-biased manner, while most boosting factors were not (Fig. 3h-i, Extended Fig. E6d). Exceptions from this pattern were Oct4, which we categorized as a basal early factor, but did not respond to X dosage, and Otx2, a late boosting factor, which was more highly expressed in XX than in XO cells, a trend that was however not confirmed in a different data set (Fig. 3i, Extended Fig. E6e). Taken together, we found that Xist activators are transiently upregulated at the onset of XCI. While a group of early basal activators integrates information on X-chromosome number and differentiation, a second set of later boosting factors only respond to differentiation cues.

### Reporter screens uncover functional TF-RE interactions at the *Xist* locus

In the next step, we aimed to dissect how the REs within the *Xist* locus responded to the TFs we identified. We built upon a comprehensive set of REs that regulate Xist, which we had previously identified through a non-coding CRISPRi screen^15^. To understand their regulation, we aimed to link them to the identified TFs in high-throughput. To this end, we developed a new CRISPR screen variant, where a fluorescent RE-reporter construct is used as screen readout.

In the first step, we established reporter cell lines for 21 Xist-controlling REs and a control without RE (noRE) (Fig. 4a-b, Extended Fig. E7a)^15^. These included 11 activating REs located either close to the Xist promoter (proximal) or up to several 100 kb upstream (distal), and 8 repressive elements, mostly within Xist’s antisense transcript Tsix. For each reporter line, we integrated GFP under control of the selected RE and the *Fgf4* minimal promoter into the genome of female mESCs through lentiviral transduction^52^. To verify that the reporter lines recapitulated the activity of the endogenous REs, they were differentiated for 4 days by 2iL withdrawal and the reporter activity was quantified by flow cytometry relative to the noRE control (Fig. 4c, Extended Fig. E7b-c). At day 2 and 4 of differentiation, reporter activity of Xist-activating REs correlated well with the activity of the endogenous elements, as assessed by ATAC-seq and H3K27ac CUT&Tag (Pearson correlation coefficient, r=0.2-0.6, Extended Fig. E7d-e)^15^. In contrast, correlation was low in the naive state and for Xist-repressing REs. Thus, we focussed on Xist-activating REs, which generally exhibited a stronger reporter signal and mirrored endogenous activity patterns more faithfully. These contained 4 promoter-proximal REs, including the *Xist* TSS RE58 and three segments of the RE57 element within Xist’s first exon (RE57L/M/R), as well as 7 distal REs, including the *Jpx* promoter RE61, the *Ftx*-associated RE85, the *Xert*-related REs 93, 95-97 and the *Rnf12* promoter RE127 (Fig. 4b-c)^15^. Similar to the endogenous locus, the proximal REs were already active prior to differentiation, while the activity of the distal REs increased throughout the time course (Fig. 4c).

**Figure 4:**
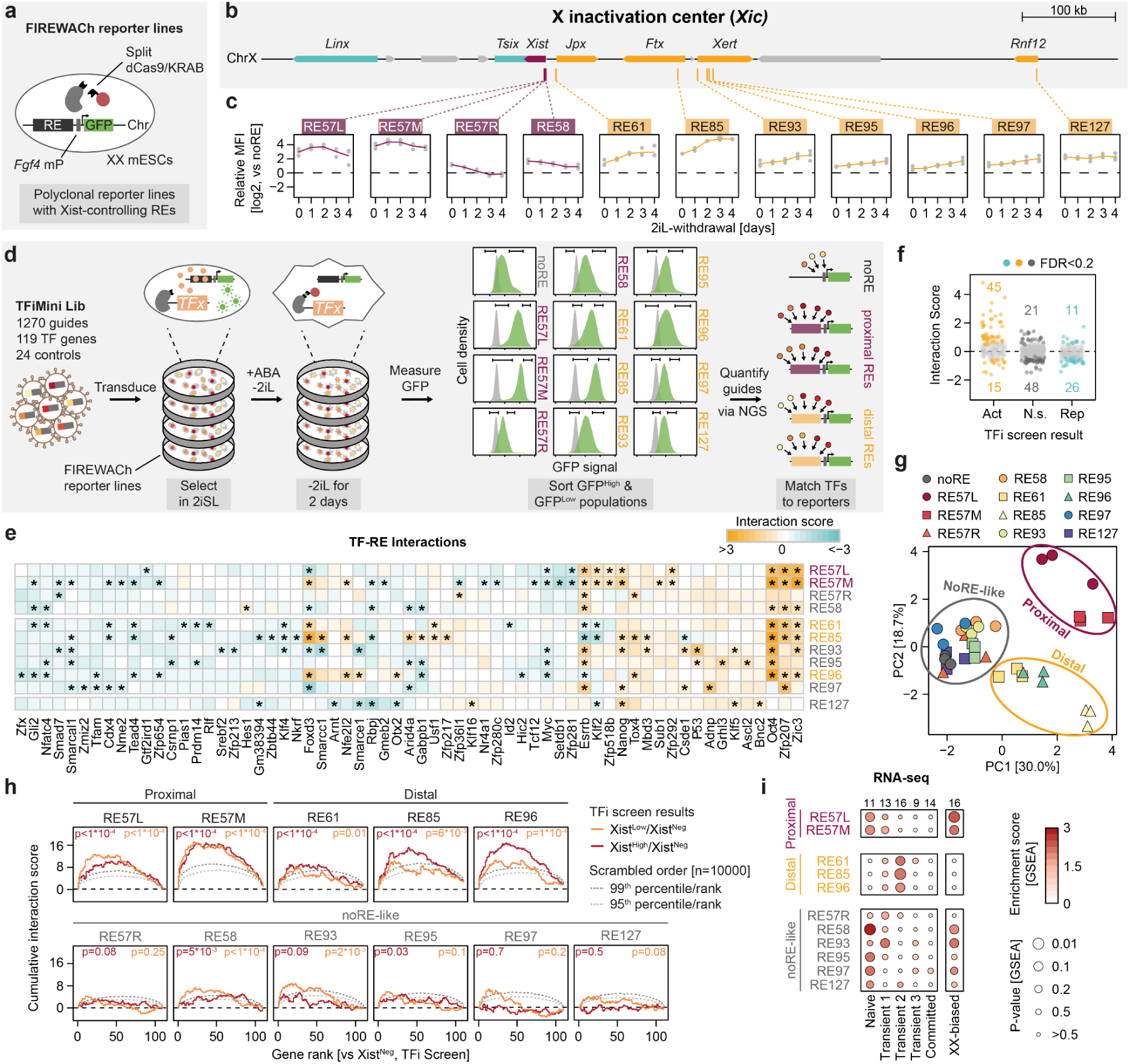
Reporter screens identify functional TF-RE interactions. (a) Schematic outline of the genomically integrated FIREWACh reporter constructs^52^. (b) The genomic landscape around the *Xist* locus with Xist regulators shown in teal (repressive) or orange (activating). Xist-controlling REs selected for the reporter screens are shown as burgundy (proximal REs) and orange boxes (distal REs). (c) Reporter expression dynamics for selected proximal (burgundy) and distal REs (orange). Relative MFI was calculated as the log_2_ fold change between GFP levels in the indicated reporter line and the noRE control. Mean (line) of n=3 biological replicates (dots) is shown. (d) Schematic outline of the reporter screens, analyzed in e-i. The TFiMini library was transduced into the 12 individual FIREWACh reporter lines. The split dCas9/KRAB system was then induced via the addition of ABA to the medium. The cells were differentiated for 2 days via 2iL-withdrawal. GFP^High^ (top 10%) and GFP^Low^ (bottom 10%) populations were sorted according to reporter expression. Guide frequencies in the different populations were quantified via NGS. (e) Heatmap displaying interaction scores between the GFP^High^ and GFP^Low^ populations for the included reporter lines. Significant interactions are indicated by a black asterisk (Benjamini-Hochberg-corrected *one-sample t-test*, FDR≤0.2). Only TF targets with at least 1 significant interaction were included in the plot. (f) Interaction scores for all assessed TF-RE combinations, with TFs grouped by effect on Xist expression in the TFi screen (Xist^High^/Xist^Neg^). Significant TF-RE interactions (FDR≤0.2) are colored in orange (Xist-activating), gray (non-significant) or teal (Xist-repressive). The number of activating (bottom) and repressive TF-RE interactions (top) per category is indicated in the plot. (g) PCA of the log_2_-transformed fold changes of all TF genes between the GFP^High^ and GFP^Low^ populations per reporter line and replicate. Groups returned by *k-means* clustering on the log_2_ fold changes are shown with thick circles. (h) Cumulative interaction scores for selected reporter screens ordered by the results of the TFi screen. TF genes are ranked from strongest activator to strongest repressor in the Xist^High^/Xist^Neg^ (red) and Xist^Low^/Xist^Neg^ (orange) comparisons. An empirical p-value, calculated by comparing the mean score of the cumulative distribution to scrambled rankings (n=10,000 bootstrap samples), is depicted in the corners. The 99^th^ (dark gray) and 95^th^ (light gray) percentiles of the bootstrap samples are shown as dotted lines. (i) Dotplot depicting GSEA results, investigating enrichment for expression groups from the RNA-seq time course in the reporter screens (left, Fig. 3). Results for a set of XX-biased factors are shown on the right (Fig.3, *ANOVA*, FDR≤0.05). Only TFs with at least one significant TF-RE interaction were included in the analysis. The number of TFs in each group is shown above the plot.

To map TF-RE wiring, we performed one CRISPRi screen per cell line (in 3 replicates) using the split dCas9/KRAB system and the TFi Mini sgRNA library. At the enrichment step, at day 2 of differentiation, we sorted the 10% cells with the highest and lowest GFP signal, respectively (Fig. 4d, Extended Fig. E8a-b). The samples generally displayed a low sgRNA distribution width and high reproducibility across replicate samples (Extended Fig. E8c-e). As positive controls, the TFiMini Lib contained sgRNAs targeting the minimal *Fgf4* promoter and the different RE inserts. These were depleted from the GFP^High^ population as expected (Extended Fig. E9a). Notably, 7 factors, including the strong Xist activator Nfrkb, significantly affected the noRE reporter, probably by regulating the minimal *Fgf4* promoter (Extended Fig. E9b), and were removed from all further analyses.

To quantify functional interactions, we defined an interaction score: We computed the z-score of the log_2_-transformed GFP^High^-to-GFP^Low^ sgRNA ratios and normalized them to the values obtained in the noRE control screen (see methods for details). In total, the analysis revealed 166 functional TF-RE interactions (FDR≤0.2, *one-sample t-test*) (Fig. 4e). The number of interactions per RE (5-27) and per TF (1-10) were highly variable (Extended Fig. E9c-d). The direction of the identified interactions (activating/repressive) typically matched the results of the TFi screen (Fig. 4f). Moreover, for TF-RE interactions involving Xist activators identified in the TFi screens, the presence of the cognate TF binding motif within the RE was associated with an increased interaction score (Extended Fig. E9e). This suggests that potent Xist activators in part act via direct binding.

We noticed that the number of identified interactions per RE correlated with signal strength of the reporter line (Extended Fig. E9f), suggesting that reporter activity might limit screen sensitivity. Also the correlation between TF and RE screen results was generally stronger for reporter lines with a strong signal (Extended Fig. E9g). Finally, principal component analysis (PCA) based on the GFP^High^-to-GFP^Low^ sgRNA ratios (Fig. 4g) revealed that only a subset of five screens, again with high reporter activity, were well separated from the noRE control, while the six remaining screens clustered close to the control (Fig. 4g). In the subsequent analyses we therefore focussed on a subset of reporter screens with good quality control metrics and high reporter activity, which formed two distinct clusters in the PCA analysis: a proximal cluster containing REs 57L/M, and a distal cluster, consisting of REs 61, 85 and 96 (Fig. 4g). All of these REs were most strongly activated by Oct4, suggesting that the TF acts as a master regulator of the *Xist* locus during random XCI (Fig. 4e). Also Zic3 broadly interacted with the indicated reporters, but most strongly with the proximal REs 57M/L. In contrast, Otx2 was specifically enriched at the distal RE96.

To assess whether the TFs interacting with specific REs play distinct functional roles in Xist regulation, we integrated the reporter screen results with the data from the TFi screens and the RNA-seq time course analysis. To compare TFi and reporter screens, we first ranked all the genes included in the reporter screens depending on their score in the Xist^Low^ (basal activation) and Xist^High^ (boosting factors) populations in the TFi screen. We then plotted a cumulative interaction score for each individual RE and compared it to the cumulative interactions score for random orders through a bootstrapping approach (Fig. 4h). Regulators of distal REs exhibited stronger interactions with factors identified in the Xist^Low^ compared to the Xist^High^ population in the screen. In contrast, proximal REs responded similarly to factors identified in the two populations. This suggests that the factors controlling distal REs tend to boost Xist levels, while those acting on proximal elements also control basal Xist activation.

In the next step, we investigated the expression patterns of the TF genes interacting with the different REs. To this end, we performed gene set enrichment analysis (GSEA) using our RNA-seq time course expression clusters to evaluate the expression dynamics of factors controlling each RE (Fig. 4i). For the proximal REs 57L/M, high interaction scores were associated with TF genes in the naive and transient 1 groups. Activating interactions for RE 61, 85 and 96, on the other hand, were enriched for TF genes in the transient 2-3 groups (Fig. 4i). GSEA using our previously defined set of XX-biased TFs (*ANOVA*, FDR≤0.05, Fig. 3f) revealed that activators of the proximal elements tended to be expressed in an XX-biased manner, while activators of the distal REs were not affected by X-chromosome number (Fig. 4i, Extended Fig. E9h-j). In conclusion, we have developed a novel methodology to probe the function of a large number of TFs at individual REs by combining traditional reporter assays with CRISPR screens. We observe that promoter-proximal regions are controlled by early factors with XX-bias, while distal enhancers within *Jpx*, *Ftx* and *Xert* tend to respond to late factors that boost Xist levels.

### Basal Xist activation and expression levels are tuned by distinct TFs

Our global analyses revealed a pattern, where basal activation appeared to be associated with early regulators acting via proximal elements, while boosting factors were upregulated later and acted via distal enhancers. To test this emerging model, we chose two early basal activators (*Nfrkb*, *Zic3*) and two late boosting factors (*Otx2*, *Foxd3*) for in-depth analysis (Fig. 5a). We created knockdown lines using our CasTuner system^41^, resulting in 80-95% knockdown and 2.5-5.5 fold reduction of Xist levels assessed by RT-qPCR (Extended Fig. E10a-c). To understand whether the regulators affect similar or distinct regulatory axes, we quantified the expression of a broader panel of Xist regulators upon knockdown (Extended Fig. E10a). The factors generally did not affect each other’s expression, except repression of *Nfrkb* following *Otx2* knockdown, and had limited effects on other genes within the X-inactivation center.

**Figure 5:**
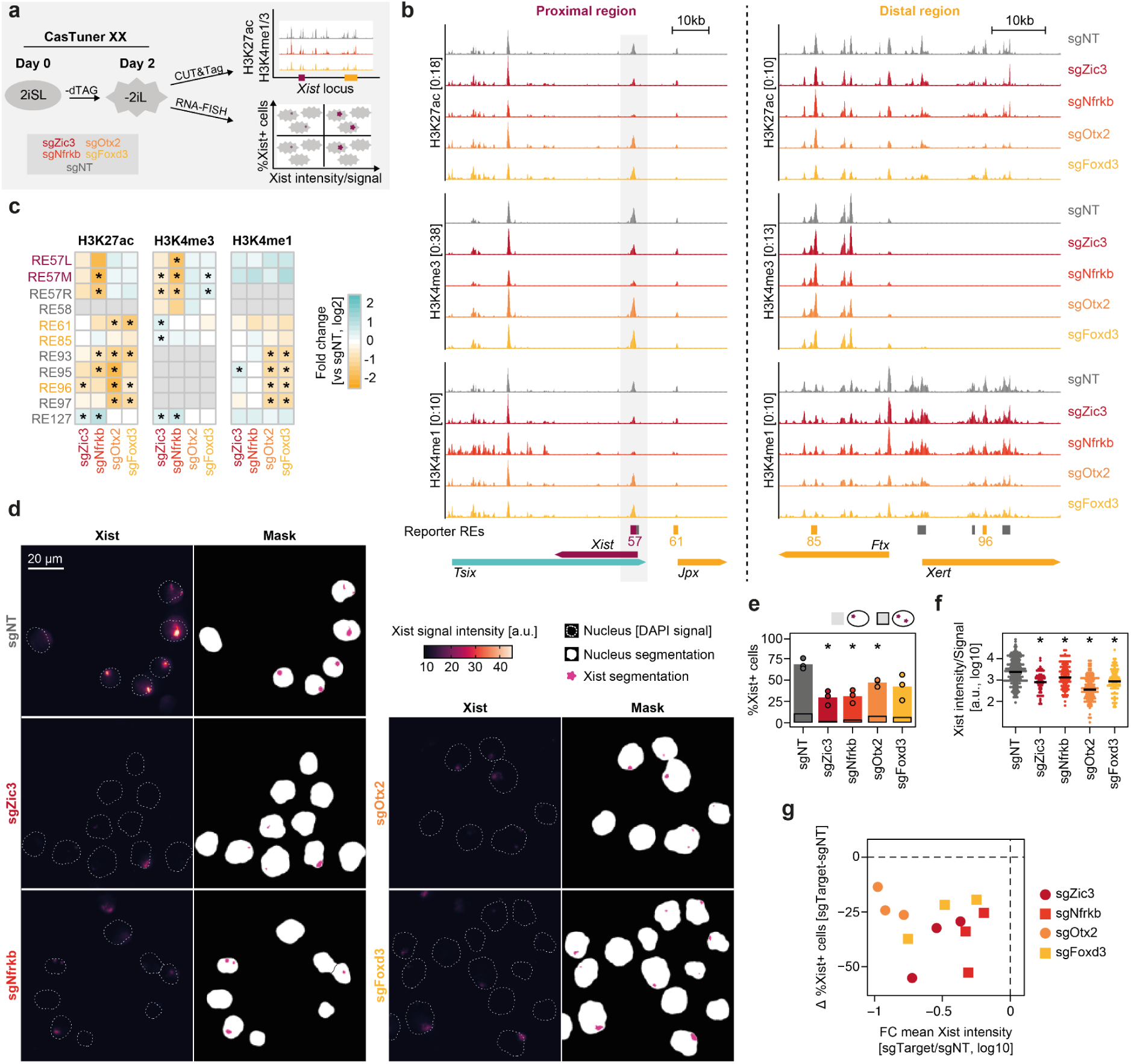
Basal Xist activation and full transcript levels are governed by distinct mechanisms. (a) Schematic outline depicting the experimental setup. CasTuner lines targeting *Zic3*, *Nfrkb*, *Otx2* and *Foxd3* were differentiated together with a non-targeting control (sgNT) for 2 days by 2iL-withdrawal. Xist RNA signals and active chromatin marks at Xist-controlling REs were assayed via RNA-FISH (d-g) and CUT&Tag (b-c), respectively. (b) Browser tracks of CUT&Tag targeting active histone marks following knockdown of different Xist activators. The promoter-proximal region around *Xist* (left) and the distal region around *Ftx* and *Xert* (right) are shown. A gray box marks the *Xist* promoter (left). The location of the reporter screen REs is shown below the tracks. Three biological replicates were performed per condition and merged for visualization. (c) Quantification of the CUT&Tag data shown in (b) at the endogenous REs assayed in the reporter screens in Fig. 4. A log_2_ fold change of the normalized counts, averaged across replicates, relative to the sgNT control is shown per RE. Significance when comparing n=3 replicates is marked by a black asterisk (*unpaired t-test*, p≤0.05). REs with <5 average counts were excluded from the analysis (in gray). (d) Exemplary RNA-FISH images depicting Xist RNA signal following the knockdown of different Xist activators. Nuclei (dotted line, white shading) and Xist signals (pink) were segmented using an automated image analysis pipeline. (e) Effect of TF gene knockdowns on the percentage of Xist+ cells. Xist signals were detected via an automated analysis pipeline. The mean (bar) of n=3 biological replicates (circles) is shown. The average percentage of cells with biallelic Xist signals is marked by a black outline. Significance versus the sgNT samples for the total percentage of Xist+ cells is marked by a black asterisk (*paired t-test*, p≤0.05). (f) Effect of TF gene knockdown on the intensity per Xist signal. Xist signals were detected and quantified via an automated analysis pipeline. The plot shows individual Xist signals from three merged biological replicates. The depicted signals were selected randomly by downsampling to match the number of the smallest replicate per condition (21-63 Xist signals per replicate sample). Significance compared to the sgNT control is marked by a black asterisk (*rank-sum Wilcoxon test*, p≤0.05). (g) Differences in Xist+ cells (from e) and mean Xist intensity (from f) following TF gene knockdown relative to the sgNT control is shown for individual replicates.

To investigate which REs are controlled by each factor at the endogenous locus, we performed chromatin profiling by CUT&Tag upon knockdown at day 2 of differentiation. We quantified H3K4me3 to estimate promoter activity, H3K4me1 as enhancer mark and H3K27ac, which generally correlates with promoter and enhancer activity^53^. While knockdown of *Zic3* and *Nfrkb* reduced H3K27ac and H3K4me3 at the *Xist* promoter, the distal REs were only weakly affected (Fig. 5b-c). Conversely, knockdown of *Foxd3* and *Otx2* had no effect on the *Xist* promoter, but induced a large reduction in H3K4me1 and H3K27ac at the distal elements (Fig. 5b-c). Interestingly, *Foxd3* and *Otx2* knockdown affected all *Xert* elements, while the reporter screens had only identified interactions with RE96 (Fig. 4e), suggesting crosstalk between REs at the endogenous locus. Nevertheless, our analysis confirmed that the early regulators Zic3 and Nfrkb indeed primarily affect proximal elements, while the late factors Otx2 and Foxd3 modulate distal regions.

To test whether the selected factors play functionally distinct roles in Xist regulation, we assessed Xist expression at the single-cell level through RNA-FISH at day 2 of differentiation. Through automated image analysis we quantified the fraction of Xist-expressing cells (frequency) and the amount of Xist RNA per chromosome (intensity). While all four factors reduced both frequency and intensity, the relative contributions of the two modalities to the total knockdown effect was variable (Fig. 5d-g, Extended Fig. E10d). The early factors had a more pronounced effect on frequency compared to the late factors, confirming a role in basal Xist activation (Fig. 5e,g). Signal intensity was affected most strongly by the late factor *Otx2*, whereas the knockdown of the early factor *Nfrkb* had minimal effects (Fig. 5f,g).

These analyses suggest that basal Xist activation and high expression levels can be controlled independently from each other. Basal activators control initial upregulation, potentially dependent on X chromosome number. Subsequently, high expression levels are obtained by activation of distal REs within *Jpx*, *Ftx* and *Xert* by a second wave of late TFs, including OTX2 and FOXD3.

### Distal REs are required for efficient silencing during random XCI

Our analysis suggests a two-step process, where basal Xist activation is distinctly regulated from high expression levels. We speculated that high Xist levels might be required for complete Xist-mediated gene silencing. To test this hypothesis we used a homozygous Δ*Ftx*-*Xert* deletion line, which lacks most of the distal REs^15^. In this mutant line, the percentage of Xist+ cells is only slightly reduced compared to wildtype cells, whereas total Xist levels are ∼2-3-fold diminished^15^. We differentiated the Δ*Ftx*-*Xert* deletion line alongside the wildtype control and performed scRNA-seq to quantify chromosome-wide Xist-mediated silencing specifically in Xist-expressing cells (Fig. 6a). We assessed day 4 of differentiation, a time point at which, unlike at day 2 used in our screens, the majority of cells are undergoing XCI^36^. Since the mutant was generated in a mESC line derived from a cross between distantly related mouse strains (B6xCast), we could use allele-specific read-mapping to assess XCI in individual cells. After filtering, 502 cells across two replicates remained for analysis. We were able to analyze a median of 93 allelic counts per cell across a median of 55 X-chromosomal genes (Extended Fig. E1ab-c). The transcriptomes of the mutant and wildtype cells overlapped strongly in PCA space, suggesting no major impact of the deletion on the cellular state (Fig. 6b).

**Figure 6:**
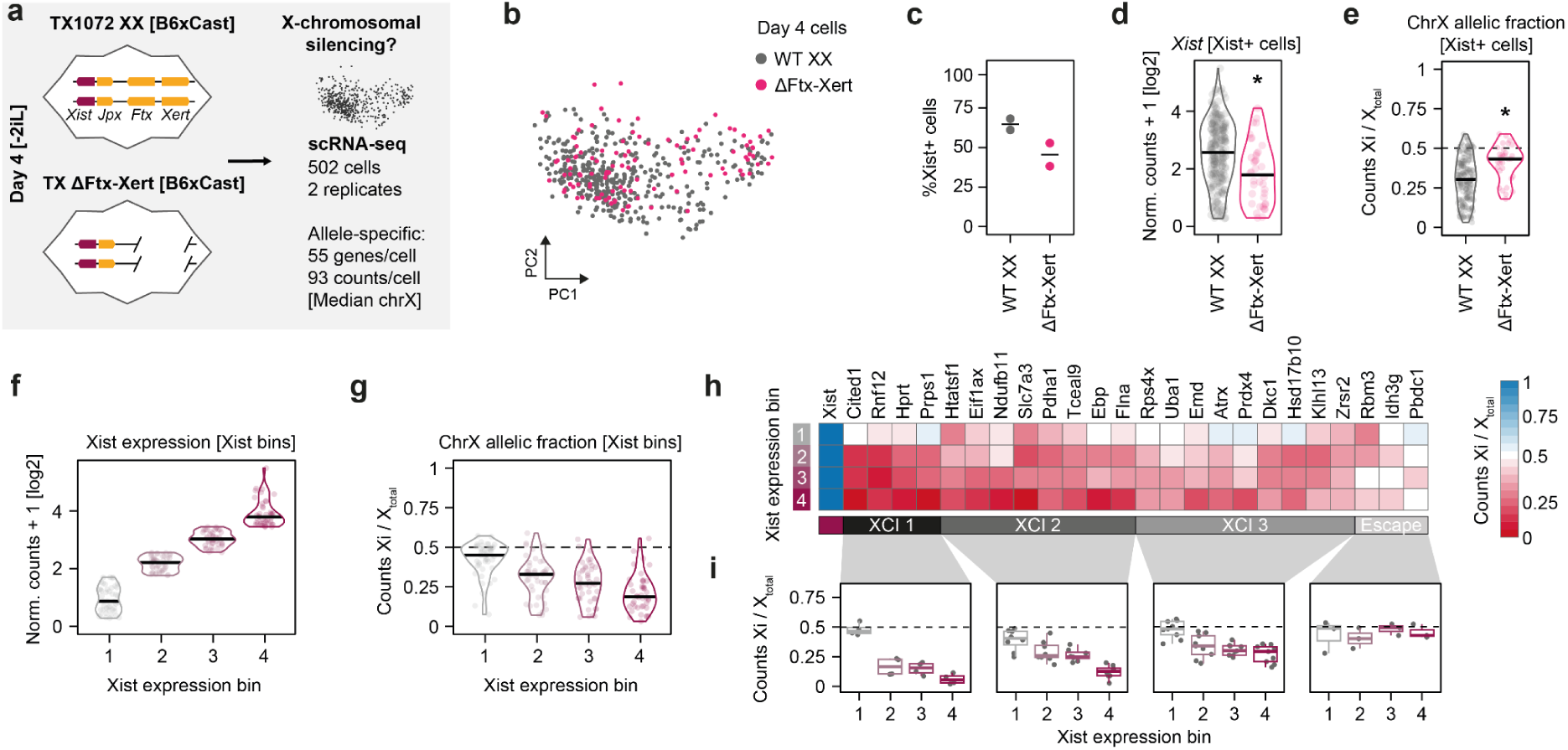
Distal REs are required for efficient silencing during random XCI. (a) Schematic depicting the Δ*Ftx*-*Xert* deletion line and the experimental outline of the scRNA-seq assay analyzed in b-i. (b) PCA depicting the transcriptomes of wildtype and Δ*Ftx*-*Xert* cells. (c) Percentage of Xist+ cells (>0 counts). The mean (line) of n=2 replicates (dots) is shown. Significance was assessed using an *unpaired t-test* (p≤0.05). (d) Normalized Xist counts per Xist+ cell (250 WT cells, 49 Δ*Ftx-Xert* cells). The replicates were merged and significance was assessed using a *rank-sum Wilcoxon test* (p≤0.05). The median is depicted by a black line. (e) ChrX allelic fraction for all Xist+ cells, quantified as the fraction of X-chromosomal allelic couts coming from the Xi. The Xi was determined as the allele with more Xist counts. Biallelic (Xist allelic fraction 0.2-0.8) and misassigned cells (chrX allelic fraction≥0.6) were excluded from the analysis (88/299 cells). Significance was assessed using a *rank-sum Wilcoxon test* (p≤0.05). The median is depicted by a black line. (f) Normalized Xist counts across Xist expression bins. Only wildtype Xist+ cells were included in the analysis in f-i (178 cells). (g) As (e), but for Xist expression bins. (h) ChrX allelic fraction across Xist expression bins for individual genes. Only genes with allelic counts in ≥15 cells per expression bin were included in the analysis (25 genes). Genes were separated into XCI groups using hierarchical clustering. (i) Data from (h) shown as boxplots. The central line represents the median and the edges of the box represent the range from the 25^th^ to the 75^th^ percentiles. Individual genes are shown as gray dots.

As expected, the fraction of Xist+ cells was slightly reduced in the Δ*Ftx*-*Xert* deletion line (63.5% vs 45.4%) (Fig. 6c), while Xist levels within Xist+ cells were strongly diminished (FC of the median = 2.01) (Fig. 6d). Next, we identified the inactive X (Xi) in each cell based on allelic Xist counts (see methods for details) and calculated the fraction of reads arising from the Xi (chrX allelic fraction), as a measure of XCI progression (Fig. 6e, Extended Fig. E11d). Compared to wildtype cells, silencing was significantly attenuated in the deletion line (Median 0.43 vs 0.30, Fig. 6e). Silencing was reduced across all X-chromosomal genes (Extended Fig. E11e). This result confirms that high Xist levels driven by distal enhancers are indeed required for efficient inactivation of the X chromosome.

To investigate the relationship between Xist levels and XCI progression in more detail, we grouped wildtype cells according to Xist expression into four bins (Fig. 6f). X chromosome-wide silencing was generally enhanced with increased Xist levels (Fig. 6g, Extended Fig. E11f). We further inspected allelic fractions at the level of individual X-chromosomal genes and performed hierarchical clustering to identify groups with similar dependency on Xist levels for silencing (Fig. 6h-i). To obtain high-confidence groups, we limited our analysis to genes that were detected in an allele-specific manner in a majority of cells (25 genes). We characterized these gene groups with respect to their genomic location, expression level (in XO cells) and silencing speed, which we had estimated in a previous study (Extended Fig. E11g-i)^54^. Interestingly, we detected a group of genes (XCI1) that displayed almost complete silencing in the second lowest Xist expression bin. The 4 genes in the group, including the Xist activator *Rnf12*, were located close to the *Xist* locus and silenced rapidly (Fig. 6h-i, Extended Fig. E11g-i). In contrast, the two other groups of silenced genes (XCI2/3) were more sensitive to Xist levels and only efficiently silenced in cells with higher Xist expression (Fig. 6h-i). Using the same gene groups, we detected a similar pattern using a different data set from a previously published scRNA-seq experiment (Extended Fig. E11j)^36^. This analysis suggests that a subset of X-linked genes, including *Rnf12*, are silenced already at rather low Xist levels, while efficient silencing of the entire X chromosome might require higher amounts of Xist RNA driven by boosting factors, such as OTX2 and FOXD3, acting via distal REs in the *Ftx/Xert* region.

## Discussion

Here, we use a combination of pooled CRISPR screens to systematically characterize input-decoding by an entire developmental gene locus. First, we comprehensively identify the *trans*-regulators of Xist at the onset of random XCI using traditional CRISPRi screens and detect a set of previously unknown regulators. We then link the identified TFs to Xist-controlling REs through several CRISPR screens with fluorescent RE reporters as the phenotypic readout. By integrating the screen results with TF gene expression patterns, we show that Xist is activated during formative pluripotency in a two-step process, where two functionally distinct groups of Xist regulators are sensed by different RE classes: While high Xist levels are driven by distal enhancers, initial Xist upregulation is governed by a set of XX-biased factors that primarily control promoter-proximal REs.

Several of the previously unknown Xist activators we have identified, including ZIC3, OTX2, FOXD3 and RBPJ, have been previously implicated in the exit from the pluripotent state^42,43,55,56^. This suggests a tight coupling of this developmental transition and XCI establishment. A subset of factors is expressed in an XX-biased manner prior to initial Xist activation, suggesting involvement in the regulation of female specificity of Xist expression. These basal activators primarily control promoter-proximal REs, which enables binary control of the gene. One of these factors is ZIC3, a TF encoded on the X chromosome. Similarly to RNF12, ZIC3 might act as a dose-sensitive X-linked Xist activator^19,28^. The remaining XX-biased factors identied in our study are encoded on autosomes. How X-chromosome number regulates these factors remains unclear. Given that only a subset of target regions respond to small changes in TF abundance^57,58^, it remains to be tested whether their observed ∼2 fold higher expression in XX compared to XO cells indeed drives XX-specific Xist upregulation.

In addition to the basal XX-biased activators, we describe a group of factors required to maximize Xist expression, which we term boosting factors. They preferentially interact with the distal Xist REs within *Jpx*, *Ftx* and *Xert*, which we have previously shown to be insensitive to X-chromosome number^15^. Accordingly, their expression levels are unaffected by the number of X chromosomes in the cell. Boosting factors are mostly dispensable for basal Xist activation, but drive high expression levels to guarantee efficient establishment of X inactivation. Similarly, it has recently been shown that Xist RNA abundance must be tightly controlled to prevent ectopic escape of X-linked genes in somatic cells and erroneous silencing of autosomal genes^59–61^.

In our proposed model, Xist upregulation occurs in two distinct stages, each controlled by different activator groups. This model parallels a previous study, where two transcription factors at different developmental stages work together to ensure lineage-specific expression of the *lsy-6* gene in C. elegans^62^. An initial “priming” step by the early activator keeps the locus accessible and sensitive to a later regulator that drives high transcript levels. This mirrors our finding that basal activators primarily promote an active chromatin state of the Xist promoter region. One such activator, NFRKB, like the established Xist activator YY1, functions as a subunit of the INO80 chromatin remodeling complex^24,44^, likely maintaining promoter accessibility. A similar association of proximal elements with binary gene control was described for the *Rex1* gene in mice^63^. It is expressed in pluripotent cells, but later silenced by promoter DNA methylation, making it insensitive to distal REs in the same topologically associating domain. Another example is the *Pitx1* locus during limb development, where expression is reduced by 35-50% upon deletion of the distal enhancer *Pen,* while the number of PITX1-positive cells are only decreases by 8-17%, supporting a primary role for *Pen* in boosting expression levels^64,65^. The principle that basal activation and transcriptional boosting are mediated by proximal and distal regulatory regions, respectively, may thus broadly apply to developmental gene regulation.

Our study describes a novel approach to investigate developmental gene regulation in a locus-wide fashion. Our reporter screens present a valuable tool for assaying single loci, where the number of regions of interest is limited. We assay 1210 TF-RE combinations and detect a total of 166 interactions. Importantly, these regulatory links represent functional interactions and not correlations or TF binding. The in-depth characterization of a single *cis*-regulatory landscape also enables a direct comparison of RE characteristics within the endogenous locus and within an ectopically inserted reporter cassette. At the *Xert* locus TF interactions diverge between the two conditions. In the endogenous context all four associated REs (93,95-97) exhibit reduced active chromatin marks upon knockdown of *Otx2* or *Foxd3*, while only RE96 is regulated by these factors at the ectopic site. One interpretation could be that the *Xert* locus is regulated hierarchically with RE96 being necessary for the activation of the other enhancers. This regulatory relationship is reminiscent of the *Pen* enhancer at the *Pitx1* locus and of facilitator elements recently described at the *α-globin* locus, which can potentiate nearby enhancers^64,66^. Similarly, RE96 displays comparable reporter strength to REs 93, 95 and 97, but has a greater impact on Xist expression upon perturbation at the endogenous locus^15^.

A possible approach to further scale up TF-RE mapping is the employment of amplicon RNA-seq as phenotypic readout instead of reporter fluorescence. While the first steps towards that goal have recently been made, the sensitivity and reliability of this approach remains to be investigated^67^. It might also overcome the challenge to identify regulators of REs that drive only low reporter activity. In our assay, the number of identified interactions increased with the reporter strength. It remains unclear whether this observation stems from technical limitations or from inherent biological differences, where stronger REs are generally regulated by more TFs. Finally, our approach identifies functional interactions, but cannot easily distinguish direct from indirect effects for individual TF-RE links. This limitation could be alleviated by integrating information on TF binding. Recent studies have developed a massively parallel binding assay (MPBA) and a multiplexed version of ChIP-seq (ChIP-DIP) that might fill that gap in the future^12,68^.

## Acknowledgements

We would like to thank the FACS facility, the microscope facility, the sequencing facility and the IT service at the MPI for Molecular Genetics for support. We thank Jana Tünnermann and René Buschow for help with setting up the automated FISH analysis. This work was supported by the Max-Planck Lise Meitner Excellence program, ERC Starting Grant CisTune (948771) and by the DFG priority program SPP 2202 (508000619) to E.G.S. T.S. was supported by the DFG (GRK1772, IRTG 2403). G.N. was supported by the European Union’s Horizon 2020 Research and Innovation Program (Marie Skłodowska-Curie ITN PEP-NET).

## Author contributions

TS and EGS conceived the project. GN generated the TX1072 XX SP427 cell line and performed RNA-seq. EK performed CUT&Tag. JJF performed scRNA-seq. MB and JJF processed scRNA-seq data. AA integrated *in vitro* RNA-seq with *in vivo* scRNA-seq data. JS performed RA and EpiLC differentiations. TS and VF generated FIREWACh reporter plasmids and performed reporter time course experiments. TS performed all other experiments and analyses. TS and EGS wrote the manuscript. Funding acquisition by EGS and MV.

## Data availability

Sequencing data generated in this study is available on GEO under accession number GSE274507. RNA-FISH microscope images are available at doi.org/10.5281/zenodo.12821363 (Related to Fig. 2, Oct4 KDs) and doi.org/10.5281/zenodo.12821095 (Related to Fig. 5, Xist activator KDs). Flow cytometry data (.fcs files) is available at doi.org/10.5281/zenodo.12822424 (Related to Figs. 1 and 4).

## Code availability

Code used in this study is available at https://github.com/EddaSchulz/TFiScreen_Paper.

## Experimental Methods

### Cell lines

The female mESC line TX1072 (clone A3) is an F1 hybrid cross between C57BL/6 (B6) and CAST/EiJ (Cast) mouse strains. It carries a doxycycline-inducible promoter in front of the *Xist* gene on the B6 allele and an rtTA insertion at the *Rosa26* locus^50^. The TX1072 XO line (clone B7) has lost the B6 X chromosome and is trisomic for chromosome 16^36^. The TX Δ*Ftx-Xert* line (clone F9) carries a homozygous deletion spanning the activating REs in *Ftx* and *Xert* (RE85-RE97)^15^. The female TX SP107 cell line (clone B6) stably expresses a split CRISPRi system, consisting of PYL1-KRAB-IRES-Blast and ABI-tagBFP-SpdCas9. Dimerization of the PYL1 and ABI domains was induced via addition of 100 µM ABA to the cell culture medium 24 hours before differentiation^15^. The female TX SP427 (clone B2) carries the CasTuner CRISPRi system, based on an HDAC4 domain fused with the conditional degron domain FKBP12^F36V^. The cell line was generated via PiggyBac-mediated transposition of the TX1072 A3 line with the pSLPB2B-FKBP12_F36V-hHDAC4-SpdCas9-tagBFP-PGK-Blast plasmid (Addgene 187956)^41^. To ensure continuous degradation of the dCas9-HDAC4 fusion protein, the cell line was cultured in the presence of 500 nM dTAG-13 (Tocris). The system was induced via removal of dTAG-13 from the medium 48 hours before differentiation. The Lenti-X 293T cell line (Takara) was used for lentiviral packaging. All ESC lines were karyotyped by shallow DNA sequencing, which confirmed a correct karyotype unless otherwise specified.

### mESC culture and differentiation

TX1072 mESCs and derived mutant lines, were cultured on 0.1% gelatin-coated flasks in serum-containing medium supplemented with 2i and LIF (2iSL conditions) (DMEM (Sigma), 15% ESC-grade FBS (GIBCO), 0.1 mM b-mercaptoethanol, 1000 U/ml LIF (Millipore), 3 mM Gsk3 inhibitor CT-99021, 1 mM MEK inhibitor PD0325901 (Axon)). The cells were cultured at a density of 4*10^4^ cells/cm^2^ and passaged every second day. If not stated otherwise, differentiation was induced at a density of 2.1*10^4^ cells/cm^2^ on 10 µg/ml fibronectin (Merck or Corning) via 2i/LIF withdrawal (-2iL conditions) (DMEM, 10% FBS (GIBCO), 0.1 mM b-mercaptoethanol). For the CRISPR screens, differentiation was induced at a density of 3.6*10^4^ cells/cm^2^. For RA differentiation, the medium was additionally supplemented with 1 µM all-trans retinoic acid (Thermo Fisher). To differentiate towards EpiLC fate, cells were cultured in serum-free N2B27 medium (50% neurobasal medium (Gibco), 50% DMEM/F12 (Gibco), 1x GlutaMAX (Thermo Fisher), 0.1 mM β-mercaptoethanol (Sigma), 0.5x N2 (Thermo Fisher), 0.5x B27 (Thermo Fisher)), supplemented with 20 ng/ml Activin A (StemCell) and 12 ng/ml FGF2 (StemCell). RNA-FISH using intronic probes of the X-linked gene *Huwe1* was regularly performed before experiments to confirm that cells had retained two X chromosomes.

### Cloning multiguide plasmids

For individual CRISPRi experiments, three different guides targeting the same promoter/RE (Supplemental Table S6) were introduced into an sgRNA expression plasmid (SP199) via Golden Gate cloning, as described previously^69^. Following cloning, each guide was controlled by a different Pol III promoter (hu6, mu6, hH1). To this end, gene blocks carrying mu6 or hH1 promoter sequences fused to an optimized sgRNA constant region (IDT) were amplified with primers containing part of the guide sequences and a BsmBI restriction site (Supplemental Table S6)^70^. The PCR-amplified fragments were then ligated into the *BsmBI*-digested (New England Biolabs) SP199 in an equimolar ratio in a Golden Gate reaction using T4 ligase (New England Biolabs) and the *BsmBI* isoschizomer *Esp3I* (New England Biolabs) for 20 cycles (5 min 37 °C, 5 min 20 °C) with a final denaturation step at 65 °C for 20 min. Vectors were transformed into NEB Stable competent *E. coli*. Successful assembly was verified by *ApaI* digest (New England Biolabs) and Sanger sequencing.

### Generation of reporter lines

The lentiviral FIREWACh system was used to create polyclonal GFP reporter lines of Xist-controlling RE, as described previously^40,52^. To this end, RE inserts were amplified from genomic DNA or bacterial artificial chromosomes (Supplemental Table S6). They were then inserted into the *BamHI*-digested (New England Biolabs) FpG5 vector (Addgene 69443) using the InFusion HD Cloning Kit (Takara)^52^. Vectors were transformed into NEB Stable competent *E. coli*. Successful cloning was verified by Sanger sequencing. The FIREWACh plasmids were integrated into the TX SP107 cell line using lentiviral transduction. Successful integration was verified using flow cytometry. A complete description of the generated lines is shown in Supplemental Table S6.

### Lentiviral transduction

CRISPR sgRNA expression vectors and FIRWACh reporter constructs were stably integrated into mESCs using lentiviral transduction. To this end, Lenti-X 293T cells were seeded at a density of 1.04*10^5^ cells/cm^2^ and transfected with a third generation transfer system consisting of VSVG, pLP1 and pLP2 (Thermo Fisher). For a 6-well plate, 1.2 µg PLP1, 0.6 µg pLP2 and 0.4 µg VSVG were incubated in 250 µl OptiMEM (Thermo Fisher) with 2 µg of the transfer plasmid. After 5 minutes, 11.25 µl Lipofectamine 2000 (Thermo Fisher), concomitantly incubated in 250 µl OptiMEM), was added. Following another incubation of 15 minutes, the mixture was transferred onto the Lenti-X cells. Viral supernatant was harvested after 48 hours and concentrated 1:10 using the Lenti-X concentrator (Takara). Concentrated supernatant was resuspended in PBS and stored at -80°C. Prior to lentiviral transduction, mESCs were cultured in SL conditions in order to prevent loss of the second X chromosome^15^. For transduction, cells were seeded at a density of 2.9*10^4^ cells/cm^2^. The next day, 8 ng/µl polybrene (Merck) was added to the culture medium to enhance efficiency. 50-200 µl concentrated virus was used per transduction. Starting 48 hours afterwards, successful integrations were enriched using antibiotic selection with puromycin (1 µg/ml, Sigma) or hygromycin B (0.2 mg/ml, Sigma or VWR). Unless indicated otherwise, cells were transferred to 2iSL medium and cultured for at least 5 passages prior to further experiments. XX-status was assessed using RNA-FISH with probes targeting the introns of the X-linked gene *Huwe1*.

### Generation of TX CasTuner line

The TX SP427 cell line (clone B2), stably expressing the FKBP-dCas9-HDAC4 CasTuner system, was generated via PiggyBac transposition^41^. TX1072 A3 mESCs were reverse transfected using lipofectamine 3000 (Invitrogen) with the pSLPB2B-FKBP12_F36V-hHDAC4-SpdCas9-tagBFP-PGK-Blast plasmid (Addgene 187956) and a vector carrying a hyperactive PiggyBac transposase (pBROAD3-hyPBase-IRES-zeocin) at a 5:1 molar ratio. Successful integrations were selected using blasticidin (5 µg/µl, Roth). The monoclonal B2 line was generated by FACS-sorting based on tagBFP expression and NGS karyotyped to ensure a normal karyotype (40, XX).

### Real-time qPCR

Cells were lysed directly on the tissue culture plates using Trizol (Invitrogen). Next, RNA was extracted using Direct-zol RNA purification Kit (Zymo Research). Subsequently, RNA was reverse transcribed into cDNA using Superscript III Reverse Transcriptase (Invitrogen) with random hexamer primers. Expression levels were quantified using the Quant-Studio 7 Flex Real-Time PCR machine (Thermo Fisher Scientific) with Power SYBR Green PCR Master Mix (Thermo Fisher). Primers used are listed in Supplemental Table S6.

### Single-molecule RNA-FISH

Exonic Xist RNA signals were quantified at the single-cell level using RNA-FISH. Hybridization was performed using Stellaris FISH probes (Biosearch Technologies)(Supplemental Table S6). To this end, cells were harvested using Accutase (Invitrogen) and adsorbed onto coverslips (#1.5, 1 mm) coated with Poly-L-Lysine (Sigma) for 5-10 min. The cells were then fixed with 3% paraformaldehyde in PBS for 10 min and permeabilized for 5 min in PBS containing 0.5% Triton X-100 (Sigma) and 2 mM Ribonucleoside Vanadyl complex (New England Biolabs). Coverslips were preserved for later use in 70% EtOH at -20°C. Hybridization was carried out overnight at 37°C with 250 nM of FISH probe in 50 ml Stellaris RNA FISH Hybridization Buffer (Biosearch Technologies) containing 10% formamide. Coverslips were washed twice for 30 min at 37°C with 2x SSC/10% formamide with 0.2 mg/ml DAPI (4’,6-Diamidin-2-phenylindol, Sigma) being added to the second wash. Prior to mounting with Vectashield (Biozol), coverslips were washed with 2xSSC at RT for 5 min. Images were acquired using a widefield Cell Discoverer 7 or Z1 Observer microscope (Zeiss) using a 100x objective.

### Flow cytometry

Fluorescent activity of the FIREWACh reporter lines and cells stained via FlowFISH was assayed using Flow cytometry^52^. For the reporter lines, cells were resuspended in FACS buffer (1% FBS in PBS with 1 mM EDTA, Thermo Fisher). Cells were analyzed and sorted using the FACSAria Fusion flow cytometer (BD). At least 20,000 cells were assayed per measurement. Sideward and forward area scatters were used to gate for live cells. Height and width of the forward and sideward scatters were used to discriminate doublets.

### FlowFISH CRISPR screens

#### CRISPR library cloning

Both libraries were cloned into the *BsmBI*-digested SP199 expression vector^69^. The oligo pool of the TFi Lib was amplified with the primers OG113/TS122 for 14 cycles in 7 individual PCR reactions with the KAPA HiFi PCR ReadyMix (Roche). As the oligo pool of the TFiMini Lib was ordered with another library, it was amplified in two steps. First, the oligo pool was separated from the other library using the KAPA HiFi PCR ReadyMix with OG113/LR256 for 12 cycles. Then cloning overhangs were added using 4 separate PCR reactions with 500 ng each, using the primers OG113/TS122. For both libraries, two Gibson cloning reactions were performed using 7 ng of the insert and 100 ng of SP199. The reactions were pooled, ethanol-precipitated and resuspended in 5 µl water. The eluted DNA was transformed into 20 µl MegaX DH10B electrocompetent cells (Thermo Fisher). Successful cloning was confirmed using Sanger sequencing and restriction digests with *BsmBI*/*XhoI* (New England Biolabs). The coverage of the libraries was determined as 513x (TFi Lib) and 8472x (TFiMini Lib). Furthermore, both libraries were amplified using the KAPA HiFi PCR Readymix for 12 cycles to add sequencing overhangs. The TFi Lib was sequenced 150 bp paired-end on the MiSeq platform yielding ∼1*10^7^ fragments. The TFiMini Lib was sequenced 100 bp paired-end on the NovaSeq 6000 platform yielding ∼4.3*10^7^ fragments. A low log_2_ distribution width of the guide counts of 2.2 (TFi Lib) and 1.5 (TFiMini Lib) confirmed that high coverage was retained during cloning (Extended Fig. E1h, E3f). All guides were present in both libraries.

#### Titer estimation

The cloned libraries were packaged into lentiviral particles as described above. Prior to transduction, the viral titers were estimated. To this end, the TX SP107 (TFi screen) and TX SP427 (TFiMini screen) cell lines were transducd in 6-well plates with 10-fold serial dilutions of the respective lentiviruses (10^-2^-10^-7^) in duplicates. After two days, antibiotic selection was performed using puromycin (1 µg/ml, Sigma). After 7 days, the surviving colonies were counted in the wells. The viral titers were estimated as 6.8*10^5^ TU/ml for the TFi screen and 9.4*10^5^ TU/ml for the TFiMini screen.

#### Tissue culture and cell sorting

Both screens were performed in two replicates. A coverage of >300x was retained during all steps. For the TFi screen, cells transduced with non-targeting guides or guides targeting RE57 were cultured alongside the library as controls. Following transduction of the TFi Lib into the TX SP107 cell line under SL conditions (MOI = 0.3), the cells were selected for three days using puromycin. Subsequently, the cells were transferred into 2iSL medium. At the same time, 1*10^7^ cells were frozen (Selected population). After three days in 2iSL conditions the split dCas9-KRAB system was induced via the addition of ABA. The following day, the cells were differentiated for 48 hours via 2i/L-withdrawal and harvested for analysis.

For the TFiMini screen, the TX SP427 cell line was transduced under SL conditions (MOI = 0.3) in the presence of dTAG-13. Following three days of puromycin selection, the cells were transferred to 2iSL conditions (Extended Fig. E3b). At the same time, 1*10^7^ cells were frozen (Selected population). After 8 days, the CasTuner system was induced via the removal of dTAG-13 from the medium. A flask with medium containing dTAG-13 was taken along as a control. Two days later, the cells were differentiated via 2i/LIF-withdrawal. At the same time, 1*10^7^ cells were frozen (2iSL population), to confirm the inducibility of the system. After two days of 2i/L-withdrawal, the cells were harvested for analysis.

#### FlowFISH

The PrimeFlow RNA AssayKit (Thermo Fisher) was used to assay Xist RNA. The assay was performed in conical 96-well plates with 5*10^6^ cells per well. Xist RNA was labeled with Alexa-Fluor647 using a type 1 PrimeFlow probe (VB1-14258, Thermo Fisher). Finally, cells were resuspended in PrimeFlow RNA Storage Buffer (Themo Fisher) and analyzed using flow cytometry. For the TFi screen, 1*10^7^ cells were frozen during the protocol (Unsorted population). Cells were sorted according to Xist expression into Xist^high^ (top 15%), Xist^basal^ (bottom 15% of the Xist+ cells) and Xist^neg^ (bottom 15%) populations. Xist+ cells were determined based on the 99th percentile of the Xist-signal from a 2iSL sample (cells that do not express Xist). At least 1.5*10^7^ cells were recovered per population. For the TFiMini screen, only Xist^High^ and Xist^Neg^ populations were sorted. At least 5*10^5^ cells were recovered per population.

#### DNA isolation and library preparation

Sequencing libraries were prepared from all indicated populations. To this end, genomic DNA was isolated using phenol/chloroform extraction. First, cell pellets were incubated for 14 hours at 65 °C in 250 µl de-crosslinking buffer (1% SDS (Invitrogen), 1.25 µl DTT (Roth) and 10 µl 5M NaCl (Sigma) in Tris-EDTA buffer (Sigma)). Next, 20 µl RNAse A (10 mg/ml, New England Biolabs) were added and the solution incubated for 1 hour at 37 °C. Next, 5 µl Proteinase K (20 mg/ml, Ambion) was added and the solution incubated for 1 hour at 50 °C. Subsequently, 275 µl of Phenol/Chloroform (Roth) were added and the mixture vortexed for 1 min. The samples were then centrifuged for 10 min at 13,000 rpm. The aqueous phase, containing the genomic DNA, was transferred to a new tube and the sample was cleaned using ethanol precipitation. The pellets were dried and resuspended in 50 µl water.

The guide cassette was amplified for 20 cycles using the KAPA HiFi PCR ReadyMix with primers OG115/OG116. To keep 300x coverage, at least 20 µg (TFi screen) or 1.6 µg (TFiMini screen) of genomic DNA were amplified per sample. Between 0.1-2 µg were amplified per reaction, as the PCR tended to be inhibited in samples stained via the FlowFISH protocol^15^. Subsequently, PCR reactions of the same samples were pooled and concentrated using the DNA Clean & Concentrator Kit (Zymo Research). Lastly, sequencing barcodes were added in a second PCR using the KAPA HiFi PCR ReadyMix for 11-12 cycles (for primer sequences see Supplemental Table S6). The TFi screen was sequenced 100 bp paired-end on the NextSeq 500 platform, yielding ∼2-7*10^6^ fragments per sample. The TFiMini screen was sequenced 100 bp paired-end on the NextSeq 2000 platform, yielding 4-8*10^6^ fragments per sample.

#### Reporter screens

The viral titer of the TFiMini library for the reporter screens (2.6*10^6^ TU/ml) was estimated in the TX SP107 cell line under 2iSL conditions, as described for the FlowFISH screens. The screens were performed in three replicates with >200x coverage. The TFiMini library was transduced under 2iSL conditions into the TX SP107 cell line carrying a polyclonal insertion of the FIREWACh reporter construct (see Supplemental Table S6) with one of 12 RE inserts (noRE, RE57L, RE57M, RE57R, RE58, RE61, RE85, RE93, RE95, RE96, RE97 and RE127)^52^. Following three days of puromycin selection, ABA was added to the medium in order to induce the split dCas9-KRAB system. Differentiation was induced 24 hours later using 2i/LIF-withdrawal. Prior to sorting, 2*10^6^ cells were taken per sample as the Unsorted population. Flow cytometry was used to sort cells according to their GFP expression into GFP^high^ (top 10%) and GFP^low^ (bottom 10%) populations. At least 7*10^5^ cells were recovered per population. Genomic DNA was isolated as described above for the FlowFISH screens. The guide cassette was amplified for 20-25 cycles using the KAPA HiFi PCR ReadyMix with primers OG115/OG116. At least 2.2 µg of genomic DNA were amplified per sample. Following clean-up with the DNA Clean & Concentrator Kit, sequencing barcodes were added in a second PCR using the KAPA HiFi PCR ReadyMix for 9-11 cycles (for primers see Supplemental Table S6). The samples was sequenced 100 bp paired-end on the NovaSeq 6000 platform, yielding 0.2-8*10^6^ fragments per sample.

#### polyA-enriched RNA-seq

PolyA-enriched RNA-seq was performed using the Collibri 3’ mRNA Library Prep Kit (Thermo Fisher) in TX1072 XX A3 and XO B7 cells at 0, 10, 16, 24, 30, 36, 48, 56, 72 and 96 hours of 2i/LIF withdrawal in three biological replicates. Library preparation was performed according to manufacturer’s instructions using 500 ng of RNA per sample. The amplified samples were sequenced 100 bp paired-end using the NovaSeq 6000, yielding 1.6-16*10^6^ fragments per sample.

#### scRNA-seq

ScRNA-seq was performed in two replicates at day 4 of 2i/LIF-withdrawal in a homozygous ΔFtx-Xert deletion line^15^ TX1072 XX A3 wildtype control, using the Single Cell 3ʹ Reagent Kit v3.1 (10X Genomics). The sequencing libraries were prepared together with several samples of an unrelated experiment.

For sample multiplexing, MULTI-seq barcode-lipid complexes were used according to the published protocol with minor modifications^71^. Sample barcodes were chosen from a list of compatible barcodes^72^. Lipid-modified anchor and co-anchor oligos were purchased from Sigma-Aldrich. Barcode- and library preparation oligos were purchased from Eurofins with NGSgrade quality. Per sample, 1*10^5^ cells were transferred to a 96-well ultra-low attachment plate (Costar) and kept on ice during the procedure. After two washes with PBS, the cells were incubated 5-10 min with an Anchor:Barcode solution (45 µl PBS, 5 µl Anchor:Barcode). The mixture was incubated for another 5-10 min after adding 5 µl of Co-Anchor solution. Labeling was quenched by adding 1% BSA in PBS, samples washed once in the same solution and resuspended in 0.4% BSA in PBS. Afterwards, cells were counted, pooled equally, and filtered using a Flowmi cell strainer 40 µm (Sigma-Aldrich). 2.5*10^4^ cells were used for scRNA-seq gene expression library preparation. The 10x Genomics library preparation protocol was performed according to manufacturer’s instructions (CG000315 Rev E, 10X Genomics) with the additional steps required to generate the separate MULTI-seq libraries for the sample-to-cell barcode assignment^71^. During cDNA amplification, 0.5 µL of 1.25 μM “MULTI-seq additive primer” was added. 50% of the purified gene expression cDNA was carried forward, and the cycle number in the index PCR was reduced by one accordingly, to increase library complexity^73^. During the first step of cDNA cleanup the supernatant was kept, which contained the short MULTI-seq library. This MULTI-seq library was then cleaned up two times using SPRI beads (KAPA HyperPure Beads, Roche). Sequencing adapters containing indexes (“TruSeq_RPIX_”, and “Universal_I5_with_index”) were added by PCR, using 2.5 µL primer at 10 µM each, 10 ng of cDNA, 26.25 µL Kapa HiFi HotStart ReadyMix 2x, and water to 50 µL final volume, with a program of 95 °C 5 min, 11 cycles of 98 °C 15 sec, 60 °C 30 sec, 72 °C 30 sec; followed by 72 °C 1 min and 4 °C hold. The resulting sequencing library was then cleaned up another time with SPRI beads. 10X gene expression libraries and MULTI-seq libraries were sequenced asynchronous 90/28 bp paired-end on the NovaSeq 6000 platform, yielding a minimum of 6.2*10^8^ and 4.4*10^7^ fragments, respectively.

#### CUT&Tag of histone modifications

Cleavage Under Targets and Tagmentation (CUT&Tag) was performed in three biological replicates to map active histone modifications along the genome, as described previously^74^. The assay was conducted for H3K27ac, H3K4me3 and H3K4me1 at day 2 of 2i/LIF-withdrawal in TX SP427 mESCs transduced with guides targeting different Xist activators (sgZic3, sgNfrkb, sgOtx2, sgFoxd3) or a non-targeting control (sgNT). Following dissociation with Accutase, 1*10^5^ cells per antibody were collected and quickly washed in Wash buffer (20 mM HEPES-KOH, pH 7.5, 150 mM NaCl, 0.5 mM Spermidine, 10 mM Sodium butyrate, Protease Inhibitor, and 1 mM PMSF). 10 μl Concanavalin A beads (Bangs Laboratories) were equilibrated with 100 μl binding buffer (20 mM HEPES-KOH, pH 7.5, 10 mM KCl, 1 mM CaCl_2_, 1 mM MnCl_2_) and then concentrated in 10 μl binding buffer. The cells were bound to the Concanavalin A beads by incubating for 10 min at RT on a rotator. Next, the beads were separated using a magnet and resuspended in 100 μl chilled Antibody buffer (Wash buffer with 0.05% Digitonin and 2 mM EDTA). Subsequently, 1 μl of primary antibody (Supplemental Table S6) was added and incubated on a rotator for 3 hours at 4°C. After magnetic separation, the beads were resuspended in 100 μl chilled Dig-wash buffer (Wash buffer with 0.05% Digitonin) containing 1 μl secondary antibody and incubated for 1 hour at 4°C on a rotator. The beads were washed three times with chilled Dig-wash buffer and resuspended in chilled Dig-300 buffer (20 mM HEPES-KOH, pH 7.5, 300 mM NaCl, 0.5 mM Spermidine, 0.01% Digitonin, 10 mM Sodium butyrate, 1 mM PMSF) with 1:250 diluted 3xFLAG-pA-Tn5 preloaded with mosaic-end adapters. After incubation for 1 hour at 4°C on a rotator, the beads were washed four times with chilled Dig-300 buffer and resuspended in 50 μl Tagmentation buffer (Dig-300 buffer, 10 mM MgCl_2_). Tagmentation was performed for 1 hour at 37°C and stopped by adding 2.25 μl 0.5 M EDTA, 2.75 μl 10% SDS and 0.5 μl 20 mg/mL Proteinase K and vortexing for 5 s. DNA fragments were solubilized overnight at 55 °C followed by 30 min at 70 °C to inactivate residual Proteinase K. DNA fragments were purified with the ChIP DNA Clean & Concentrator kit (Zymo Research) and eluted with 25 μl elution buffer.

Sequencing libraries were generated by amplifying the DNA fragments with barcoded primers using NEBNext HiFi 2x PCR Master Mix for 14 cycles (Supplemental Table S6). Cleanup following PCR was performed with a 1x volume of Ampure XP beads (Beckman Coulter) and samples were eluted in 27 μl 10mM Tris pH 8.0. The samples were sequenced 100 bp paired-end on the NovaSeq 6000 platform, yielding ∼4.8-7.9*10^6^ fragments per sample.

## Computational Methods

If not stated otherwise, computational analysis was performed in *Rstudio* (v4.2), primarily using the *tidyverse* packages (v1.3.0)^75,76^.

### RT-qPCR analysis

Gene expression during RT-qPCR analysis was quantified using the 2^−ΔΔCt^ method. Here, relative expression was calculated using the two housekeeping genes *Rrm2* and *Arpo*. Significance was assessed using an unpaired t-test between the indicated samples using [t.test(var.equal=TRUE)].

### RNA-FISH analysis

Xist RNA signals were either counted manually (100 cells per condition) or detected and quantified with a custom python pipeline based on the *scikit-image* toolkit (v0.22) (Fig. 5d-g)^77^. In detail, microscope images were loaded as *numpy* arrays (v1.26.2) from .czi files using *czifile* (v2019.7.2) (https://github.com/AllenCellModeling/czifile)^78^. The DAPI and Xist signals were flattened using a maximal projection along the z-axis. The nuclei were segmented using an Otsu threshold on the DAPI signals^79^. Subsequently, adjacent nuclei were separated using a watershed algorithm. Nuclei touching the image borders were removed from the analysis. In addition, nuclei were filtered depending on size and eccentricity. Subsequently, the Xist signal inside the nuclei was segmented based on the maximal projections. First, the images were treated with a gaussian blur (sigma=2). Then, Xist signals were recognized using a local Niblack algorithm with a window size of 151 pixels and a k of -3.8. Here, a threshold is set for each individual pixel, depending on the intensities of the neighboring pixels (Threshold = mean(window) - k * sd(window)). Lastly, the Xist clouds were filtered on size.

For quantification, the number of segmented signals was counted in each cell individually. Only a small minority of cells (<1%) was falsely annotated with more than two Xist signals. The intensity was quantified per Xist signal by subtracting the background intensity in each nucleus (i.e. the geometric mean of the pixels outside segmented Xist signal) from the intensity within the segmented signal. A complete list of segmented Xist signals is shown in Supplemental Table S4. Significance for the Xist signal frequency was calculated using a paired t-test with [t.test(var.equal=TRUE, paired=TRUE)]. For comparing the Xist signal intensity, the signals within each condition were downsampled to guarantee equal representation of all three replicates and replicates were merged. Significance was then assessed using a Wilcoxon rank-sum test with [wilcox.test()]. To compare relative effects of the different perturbations on signal frequency and signal intensity, the percentage of Xist-positive cells or the log_10_-geometric mean of the Xist signal intensities was subtracted from the equivalent values for the non-targeting control (Fig. 5g).

### Flow cytometry analysis

Flow cytometry data was analyzed using the *opencyto* (v.1.24.0) and *flowCore* (v1.52.1) packages^80,81^. Live cells were detected using the sideward and forward area scatter. Height and width of the forward and sideward scatters were used to discriminate singlets from doublets. Mean fluorescence intensity (MFI) was calculated as the geometric mean of the intensity in the target population minus the geometric mean of the intensity in a non-fluorescent control.

### CRISPR library design

To design a CRISPR library targeting expressed TFs during early differentiation of female mESCs (TFi Lib), a list of mouse TFs was obtained from the *AnimalTFdb3.0* (Extended Fig. E1a)^37^. Next, TF genes were selected depending on their maximal expression between day 0 and 4 of 2i/LIF-withdrawal in a published RNA-seq time course of the TX1072 A3 cell line (TPM≥10) (Extended Fig. E1b)^36^. Additionally, a set of previously identified Xist regulators was added as positive controls (Supplemental table S1). Using previously generated TT-seq data, active TSSs of these genes included in the *GENCODE M25* annotation were annotated at days 0, 2 and 4 of 2i/LIF-withdrawal^15,82^. Here, *Rsubread* (v2.0.1) was used to count reads 2 kb up- and downstream of the TSSs with [featureCounts(isPairedEnd = TRUE, strandSpecific = 2, allowMultiOverlap = TRUE)]^83^. TSSs exceeding a log_2_ fold change of 1 between the up- and downstream bins were classified as active. Nearby TSSs within 500 bp were combined into promoters (Extended Figure E1c). All promoters were then extended to 500 bp. The resulting set of 911 target promoters, including 570 TF genes and 32 non-TF controls, was used to generate the TFi library with the *GuideScan2* webtool^84^. Up to 12 guides per promoter were chosen according to their efficiency score. Three target promoters with less than three possible guides were removed from the library. 300 non-targeting guides were added as negative controls^85^. Finally, 3’ (ATCTTGTGGAAAGGACGAAACACCG) and 5’ overhangs (GTTTAAGAGCTATGCTGGAAACAGCATAGCAAGT) were added to the guide sequences. The resulting library, containing 11058 guides, was ordered as an oligo pool from GenScript.

A second CRISPR library was designed to conduct a validation screen targeting Xist and to perform the reporter screens (TFiMini Lib). All TF genes that were enriched in the Xist^High^/Xist^Neg^ comparison of the TFi screen were included (Wald.p≤0.2). Additionally, Nanog, Prdm14, Yy1 and Ctcf were added as targets, as they had been previously implicated in Xist regulation^20,24,32,86^. For each gene, the top 8 guides by absolute fold change were included, targeting the most significantly enriched or depleted promoter (by p-value). The 100 non-targeting controls with the lowest absolute fold change in the TFi screen were added as negative controls. Furthermore, the top 10 guides targeting selected REs at the *Xist* locus from a previous CRISPR screen were added to serve as positive controls for the reporter screen^15^. Lastly, we used the *CHOPCHOP* webtool to generate 10 guides targeting the *Fgf5* minimal promoter of the FIREWACh construct^52,87^. 3’ (ATCTTGTGGAAAGGACGAAACACCG) and 5’ overhangs (GTTTAAGAGCTATGCTGGAAACAGCATAGCAAGTAATGGACATCTTATTCACAG) were added to the guide sequences. The resulting library, containing 1270 guides, was ordered as an oligo pool from GenScript (Supplemental Table S1).

### FlowFISH CRISPR screen analysis

#### Data processing

Following sequencing of the TFi and TFiMini screens, the *MAGeCK* analysis toolkit (v0.5.9.3) was used to align .fastq files to the respective CRISPR libraries and generate guide count tables with [mageck count --norm-method control] (Supplemental table S1)^88^. Enrichment between different populations was calculated by using *MAGeCK* (v0.5.9.3) with the normalized count tables and [mle --permutation-round 10].

#### Quality control

Variance between the guide counts in each population was quantified as a log_2_ distribution width^89^. A high value suggests that the target coverage was lost during the protocol. Enrichment of a population should increase the distribution width slightly in the targeting guides, but not in the non-targeting controls. To this end, the normalized counts for each of the samples were first split between non-targeting controls and targeting guides. Next, the 10^th^ percentile of the counts was subtracted from the 90^th^ percentile and log_2_ transformed.

The analysis showed that a high coverage was retained in all replicates of all screens since the log_2_ distribution width remained low (∼1.5-3) (Supplemental table S1). In general, the pre-sorting fractions displayed lower distribution width than the sorted fractions. This was much less pronounced in the distribution width of the non-targeting controls. To verify the reproducibility of the screens, the Pearson correlation coefficient was calculated for each population between the different replicates. Here, the normalized counts of all guides were used with [cor()]. A summary of the published controls included in the screen is listed in Supplementary Table S1.

### RNA-seq analysis

#### Alignment and data processing

Reads were aligned single-end to the mouse reference genome (mm10) using *STAR* (v2.7.9a) with options [--outSAMattributes NH HI NM MD]^90^. One of the samples (XO_36h_R1) was removed due to very low read counts. Gene expression was quantified using the GENCODE M25 annotation supplemented with the *Xert* coordinates^15,82^. Here, *Rsubread* (v2.0.1) was used with the options [featureCounts(isPairedEnd=FALSE, GTF.featureType=”exon”, strandSpecific=1]^83^. Afterwards, counts were converted to counts per million (CPM). Mapping statistics and quality control metrics for data produced in this study are listed in Supplemental Table S2.

#### Integration of in vitro and in vivo RNAseq

To identify the exact stages corresponding to our *in vitro* setup, the RNA-seq time course data was integrated with a published *in vivo* scRNA-seq dataset of developing mouse embryos^48^ using the *scATAcat* pipeline^47^. The scRNAseq data was retrieved from GSE100597, paired-end aligned to the mm10 reference genome and reads quantified using *STAR* (v2.7.9a) with [--outSAMattributes NH HI NM MD –quantMode GeneCounts]^90^. ICM cells at E3.5, as well epiblast cells at E4.5, E5.5 and E6.5, were identified based on the expression of marker genes and summarized in pseudobulk. Subsequently, differentially expressed genes were identified across the *in vitro* bulk samples using *DESeq2* (v.1.42.0) by conditioning on genotype and time point^51^. Each time point was compared against the control (0h) and the differentially expressed genes (Wald.FDR≤0.05) were combined to determine the final gene set. These genes, serving a similar purpose to variable genes during standard scRNA-seq analysis, were used to perform PCA with the *in vitro* and *in vivo* samples. Subsequently, all samples were visualized in the same low-dimensional space using the first two principal components.

#### Clustering expression dynamics across TF genes

The RNA-seq time course experiment (Fig. 3) was used to identify groups of TF genes with similar expression dynamics during 2i/LIF-withdrawal (Supplemental Table S2). To this end, the normalized count tables were filtered for all TF genes included in the TFi screen. Next, the counts were scaled using a z-score transformation. The TF genes were sorted according to their maximal expression compared to high Xist expression into naive (0h), early (10-30h) and late factors (36-96h). Subsequently, *k-means* clustering was used with [kmeans(n = 2)] to further separate the early (into *Transient 1* and *Transient 2*) and late factors (Into *Transient 3* and *Commited*). The enrichment of Xist activators and repressors in the different groups was quantified using a one-sided Fisher’s exact test with [fisher.test(alternative=”greater”)].

Xist activators were primarily enriched in the three transient clusters (22/26). X-dosage sensitive expression dynamics of activator groups was analyzed depending on RNA-seq cluster or TFi screen results (Xist^High^ + Xist^Low^ vs Xist^High^ only). To this end, the z-scores were visualized separated by genotype using a local smooth with [geom_smooth(method=”loess”, se=TRUE)].

#### Statistical analysis

X-dosage and differentiation sensitive TF genes were determined for the entire time course using *ANOVA* with [aov(norm_counts ∼ genotype*timepoint)], followed by multiple testing correction (FDR≤0.05). Differential expression between the cell lines at individual time points was quantified using *DESeq2* (Wald.FDR≤0.05)^51^.

### Reporter time course analysis

For each sample the MFI was calculated as described above. Each RE sample was then normalized by calculating the log_2_ fold change to the noRE reporter line. This was done to control for changes in basal reporter transcription across time points. The mean MFI was then calculated as the geometric mean across replicates.

### Reporter screen analysis

Count tables were generated similarly to the Xist FlowFISH screens using *MAGeCK* (v0.5.9.3) with [mageck count –norm-method control]^88^. 39 Guides with low expression in the unsorted population (≤150 mean normalized counts) were removed from the analysis. Guides targeting *Zfp866* were removed from the analysis since their sequence overlapped with guides targeting *Gm20422*. Only guides with more than 150 mean counts in the unsorted samples were used for analysis. Detailed results are listed in Supplemental table S3.

#### Quality Control

Variance between the guide counts and reproducibility between replicates was quantified as for the FlowFISH CRISPR screens. A high coverage was retained throughout the different reporter lines and replicates (∼1.8-3.3 log_2_ distribution width). Replicate populations generally correlated well (∼0.6-0.94 Pearson correlation coefficient).

#### Analysis of positive controls

To confirm the technical success of the assay, the library design included positive control guides targeting the minimal promoter of the reporter construct or the sequence of the integrated REs. Enrichment was quantified as the log_2_ fold change between the normalized counts in the GFP^HIgh^ or GFP^Low^ populations. Guides targeting RE57 were assigned to RE57L/M/R and RE58 according to their genomic location. Only a single guide was targeted towards RE57L.

#### Analysis of the noRE reporter screen

The noRE reporter screen was analyzed separately in order to identify interactions of targeted TFs with the FIREWACh minimal promoter. Significant enrichment or depletion was detected using an unpaired t-test between the normalized counts in the GFP^High^ and GFP^Low^ populations, followed by multiple testing correction (FDR<0.2). 7 TF targets (Nfrkb, Sall4, Ctcf, Zfp296, Zfp236, Sp1 and E2f4) were removed from further analysis due to their effect on the noRE reporter line (Supplemental table S3).

#### Clustering analysis

To identify groups of reporter lines that behaved similarly during the screens, PCA was performed on the GFP^High^/GFP^Low^ log_2_ fold changes of the targeted TF genes with [prcomp(t(lfc_df))]. All non-TF controls were excluded from the analysis. The reporter lines were split into noRE-like, distal and proximal groups using *k-means* clustering on the log_2_ fold changes with [kmeans(t(lfc_df), n=3)].

#### Quantification of TF-RE interactions

Interaction scores for the assayed TF-RE interactions were calculated as z-scores across all GFP^High^/GFP^Low^ log_2_ fold changes, as done before for CRISPR screen data^91^. This was done to correct for differences in observed effect sizes between the reporter lines. In detail, mean GFPHigh/GFPLow log_2_ fold changes were calculated for all guide/reporter combinations. Each log_2_ fold change was then normalized by subtracting the mean log_2_ fold change across each target from the noRE reporter results. The corrected fold changes were scaled using a z-score transformation and named “interaction scores”. Significance was calculated using a one-sample t-test with [t.test(mu=0)], followed by multiple testing correction with [p.adjust()]. The analysis revealed a total of 166 significant TF-RE interactions (FDR≤0.2).

#### Downstream analysis

To confirm the validity of the interactions and to detect regulatory patterns, the reporter screen data was correlated to the TFi screen and RNA-seq data. First, interaction scores of the identified TF-RE interactions were compared to the Xist^High^/Xist^Neg^ comparison from the TFi screen across all (Fig. 4e) or individual reporters (Extended Fig. E9g). Similarity to the TFi screen was further assessed by ordering TF targets by their rank in the Xist^High^ or Xist^Low^ comparisons from the TFi screen and plotting a cumulative interaction score. For statistical analysis, an empirical p-value was estimated by calculating the mean cumulative score across all ranks and comparing it to the mean cumulative scores, when the ranks were scrambled (n=10,000). For visualization, the 99th and 95th percentile of the scrambled distributions at any particular rank is included in the plot. In addition, *GSEA* was performed to test for an enrichment of TF gene expression clusters (see Fig. 3) among activating interactions in the reporter screen^92^. Additionally, enrichment of TF genes sensitive to X-chromosome number was tested. To this end, the TF genes were filtered for those with at least one significant interaction in the reporter screens (FDR≤0.2) and used *fgsea* (v1.24.0) with [fgsea(minSize=5, maxSize=500, nperm=1000, scoreType=”pos”]^93^. In addition, z-score transformed RNA-seq expression of positively interacting TF genes (see Fig. 3) across individual reporter lines (Extended Fig. E9h) or distal, proximal and noRE-like reporter clusters (Extended Fig. E9i-j) was calculated for the first 48 hours of 2iL-withdrawal. XX-specific expression for the proximal and distal RE clusters was assessed using a *Wilcoxon rank-sum test* (p≤0.05).

#### Motif enrichment analysis

DNA-binding motifs of reporter screen TFs were investigated to differentiate between direct and indirect interactions. To this end, *FIMO* (v5.1.1) was used with [fimo] to identify motifs from a comprehensive set of non-redundant TF motifs. Next, the motifs were filtered for TFs with at least one functional interaction in the reporter screens. The absolute interaction scores across all REs were plotted in the different groups as a cumulative distribution depending on the presence of a TF motif. In addition, the analysis was performed after subsetting for interactions involving Xist activators (see Fig. 1). Significance was calculated using a *Wilcoxon rank-sum test* (p≤0.05).

### CUT&Tag analysis

Processing of CUT&Tag data was performed as described previously^74^. In detail, sequencing adapters were removed using *Trim Galore* (v0.6.4) with option [--paired –nextera] (https://github.com/FelixKrueger/TrimGalore). The trimmed reads were then aligned using *bowtie2* (v2.3.5.1) with the options [--local --very-sensitive-local --no-mixed --no-discordant --phred33 -I 10 -X 2000]^94^ and processed using *samtools* (v1.10) with options [view -f 2 -q 20] and [sort]^95^. Blacklisted regions for *mm10* were removed using *bedtools* (v2.29.2) with options [intersect -v]^96,97^. For the analysis of sequencing tracks with the *UCSC genome browser*^98^, .bam files of individual replicates were merged using *samtools* (v1.10) with [merge]^95^. Duplicated fragments were removed using *Picard* (v2.18.25) with options [MarkDuplicates VALIDATION_STRINGENCY=LENIENT REMOVE_DUPLICATES=TRUE] (http://broadinstitute.github.io/picard). Mapping statistics and quality control metrics are listed in Supplemental Table S4. To identify changes at *Xist*-controlling REs, reads were quantified *RSubread* (v2.0.1) with [featureCounts(isPairedEnd=TRUE]^83^. Changes compared to the sgNT control were assessed using an *unpaired t-test* (p≤0.05). RE/histone mark combinations with <5 average counts across all samples were removed from the analysis.

### scRNA-seq analysis

#### Data processing

Sequencing reads from the 10x cDNA library were mapped to the mouse mm10 genome with *STARsolo* (v.2.7.11a) with settings [soloType=CB_UMI_Simple, soloFeatures=GeneFull_Ex50pAS]^90,99^. To map reads to either the reference B6 or the alternative Cast alleles, the “Diploid Genome” options of STARsolo were used together with experimentally determined single nucleotide variants from the TX1072 cell line^54^. To remove empty droplets and assign a quality metric to the mapping of allelic reads *EmptyDrops*^100^ and *WASP*^101^ were used within *STARsolo* (v.2.7.11a)^99^. Outlier cells were filtered using default settings of *scater* (v1.26.1)^102^. MULTI-seq sample barcodes were extracted from the separately sequenced MULTIseq library and assigned to cells using *deMULTIplex2* (v1.0.1)^103^. Only cells with a uniquely assigned sample barcode were kept for further analysis.

To generate allele-specific count tables of X-chromosomal reads, reads were filtered for high-quality assignments by the presence of the *vW:i:1 WASP* filter read tag. Next, X-chromosomal reads were extracted and split into separate .bam files for each haplotype. The .bam files were sorted and indexed with *samtools* (v1.19.2)^95^. Count tables were generated using *featureCounts* (v2.0.6)^104^ to assign reads to genes and *UMI-tools* (v1.1.4) to quantify UMI-level counts with option [--per-gene --gene-tag=XT --assigned-status-tag=XS --per-cell --umi-tag=CB --cell-tag=CB –extract-umi-method=tag]^105^. Non-allelic count tables were generated in the same manner, using the unfiltered .bam file as input. Further downstream analyses were performed with *Scanpy* (v1.9.8) and *Anndata* (v0.10.5)^106,107^. Cells with ≥3% mitochondrial reads were removed. Read counts were normalized to 10,000 reads per cell and log-transformed using the [sc.pp.normalize_total] and [sc.pp.log1p] functions. To filter XO cells, cells with normalized Xist expression ≤0.5 and an X-chromosomal ratio (B6 chrX reads/total chrX reads) ≥0.85 or ≤0.15 were removed. PCA was performed using the 500 most variable genes with the functions [sc.pp.highly_variable_genes] and [sc.tl.pca].

#### Allelic analysis

To investigate X-chromosome silencing, cells with no allelic Xist counts and those with bi-allelic Xist expression were filtered out (Xist allelic fraction 0.2-0.8) (Extended Fig. E11e). In the remaining cells, the Xi was determined as the allele with the higher Xist count. The allelic fraction was then calculated for the entire chrX or individual genes with Counts_Xi_ / Counts_Xi+Xa_. Cells with misassigned Xi were removed (chrX allelic fraction≥0.6). To identify groups of genes with similar silencing dynamics, all wildtype cells were grouped into four equally sized groups according to normalized Xist expression. Next, allelic ratios were calculated for all X-chromosomal genes with ≥1 counts in ≥15 cells per expression bin. Gm14513 and Rps24-ps3 were removed from the analysis, as all allelic reads mapped to the same haplotype. The genes were classified into different groups using hierarchical clustering with [hclust(type=”ward.D2”)]. The resulting groups were further characterized depending on linear distance to the Xist TSS, expression in the absence of XCI (RNA-seq in day 4 XO cells, see Fig. 3) and estimated silencing half-time upon ectopic Xist induction^54^. To confirm the XCI dynamics in the different gene groups, allelic expression was similarly quantified using Xist+ cells (≥5 Xist UMIs) from a published scRNA-seq time course^36^. As day 4 cells had already completed XCI in this dataset, the analysis was performed using day 3 cells. Allelic counts for individual cells are listed in Supplemental table S5.

## Supplemental Material

**Extended Figure E1:**
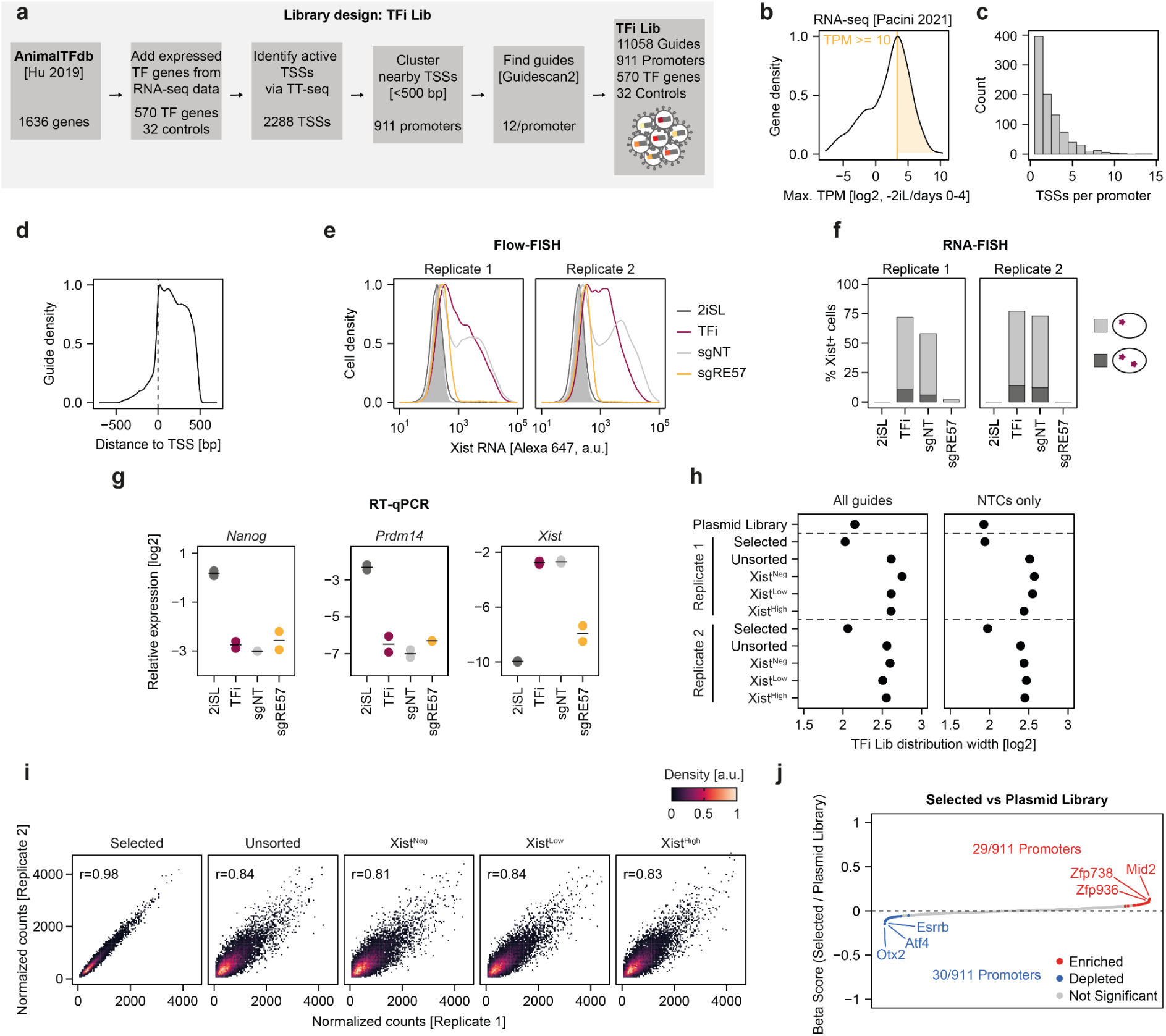
(a) Workflow to design the TFi library. Expressed TF genes during the first 4 days of 2iL-withdrawal were filtered from the *AnimalTFdb 3.0* using a published RNA-seq time course (maximal TPM>10)^36,37^. Active TSSs were then detected using published TT-seq data^15^. Nearby TSSs (≤500 bp) of the same genes were grouped into promoters. *Guidescan2* was used to generate up to 12 guides per promoter^84^. (b) Selection of expressed TF genes using a published RNA-seq time course of 2iL-withdrawal. Max TPM denotes the highest expression value between days 0-4 of differentiation. The chosen threshold of 10 TPM is indicated by the orange line. (c) Histogram depicting the number of annotated TSSs per combined promoter. (d) Density plot showing the location of the designed guides relative to annotated TSSs. (e) Xist RNA expression assessed by flow cytometry following FlowFISH staining. Additionally to the library (TFi) the plot also depicts an undifferentiated sample (2iSL), a non-targeting (sgNT) and a positive control (sgRE57). (f) Percentage of Xist+ cells assessed using RNA-FISH. The fraction of cells with biallelic Xist signals is indicated in dark gray. (g) Expression of Xist and the naive pluripotency factors *Prdm14* and *Nanog* assessed by RT-qPCR. The black bars denote the mean between two replicates (dots). (h) sgRNA distribution width of the normalized counts in the different populations of the TFi screen, grouped into targeting (left) and non-targeting guides (right). The score is given by the fold change between the 90^th^ and the 10^th^ percentile. (i) Correlation between the two replicates in the different populations of the TFi screen (11,058 guides). The Pearson correlation coefficient is indicated in the plots. (j) Ranked plot depicting enriched and depleted targets between the Unsorted and Selected populations. Significance was assessed using *MAGeCK mle* (Wald.p≤0.05).

**Extended Figure E2:**
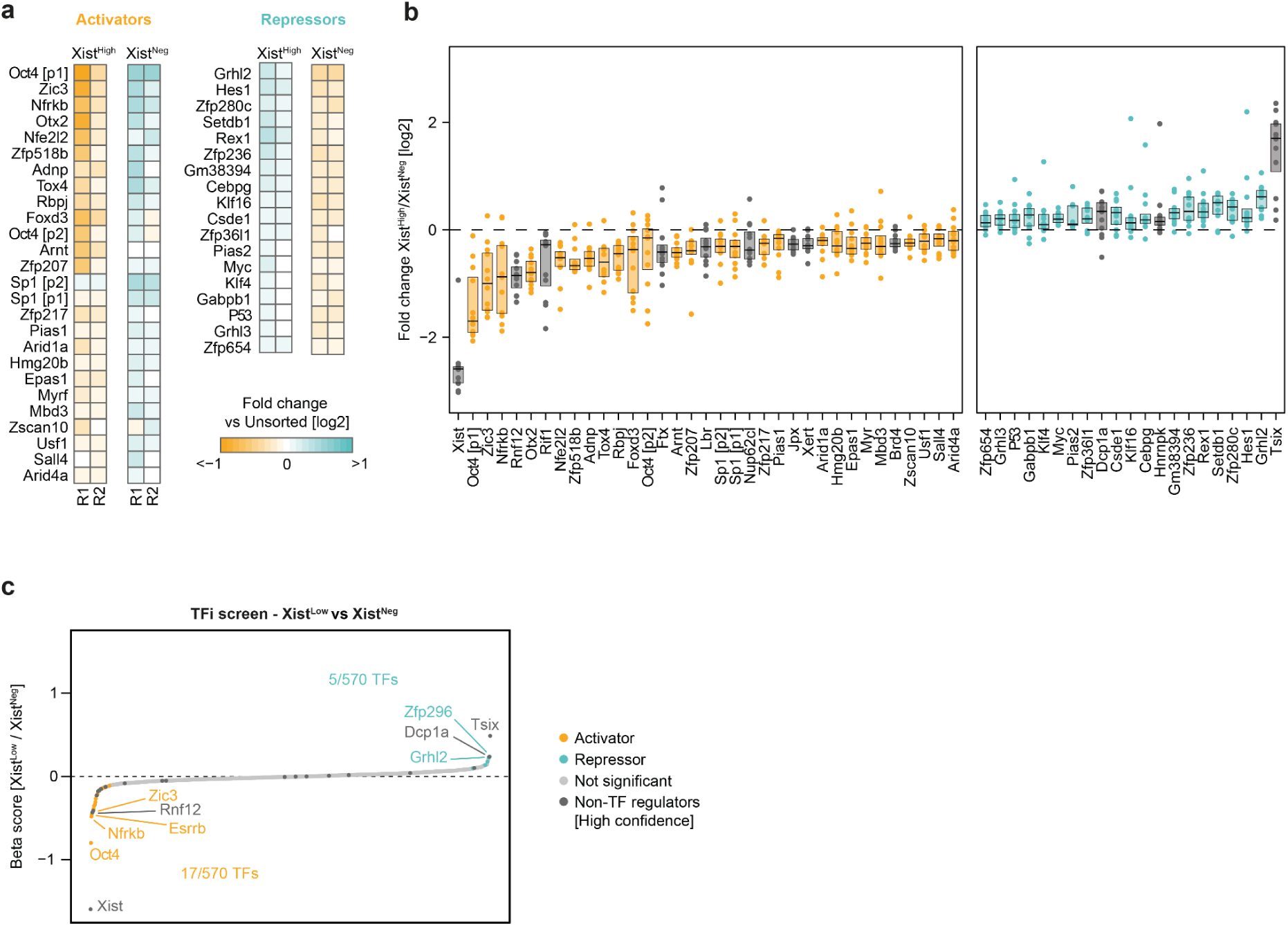
(a) Mean log_2_ fold change across guides of the Xist^High^ and Xist^Neg^ populations compared to the Unsorted cells, separated by replicates (R1/R2). All significant activators and repressors from the Xist^High^/Xist^Neg^ comparison were included in the plot (Wald.p≤0.05). (b) Log_2_ fold change of the individual guides between the Xist^High^ and Xist^Neg^ populations in the TFi screen. TF activators and repressors are colored in orange and teal, respectively. Non-TF controls are colored in gray. The boxes indicate 25^th^ and 75^th^ percentile, as well as the median (black line). (c) Comparison of target abundance between Xist^Low^ and Xist^Neg^ populations in the TFi screen. Significantly enriched or depleted targets are colored in teal or orange respectively (*MAGeCK mle*, Wald.p≤0.05). High confidence non-TF controls are colored in dark gray (see Supplemental Table S1).

**Extended Figure E3:**
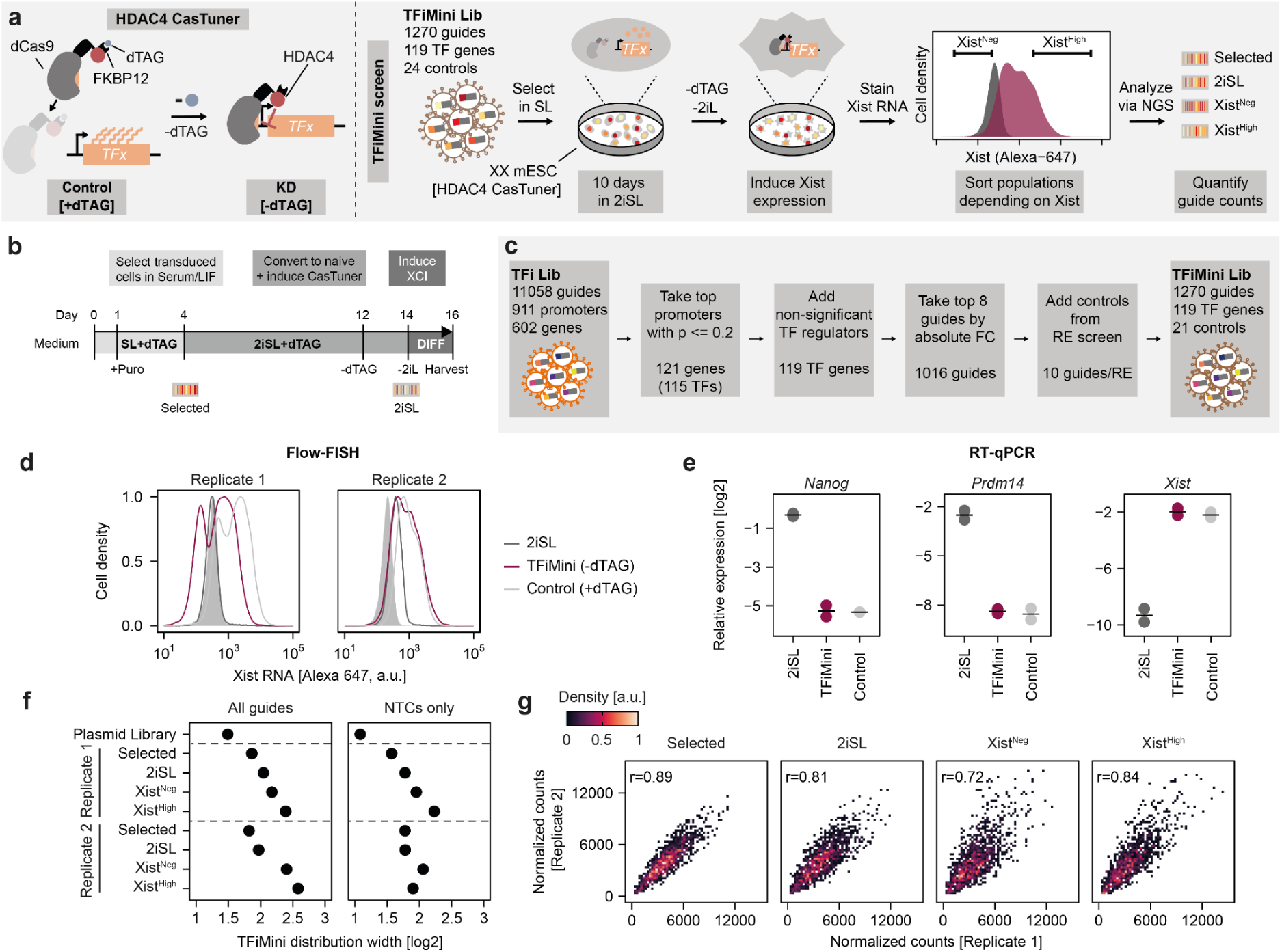
(a) Schematic of the CasTuner system (left) and the TFiMini screen (right). The TFiMini library was lentivirally transduced into TX1072 XX SP427 mESCs under SL conditions in the presence of dTAG. Following antibiotic selection and 8 days under 2iSL conditions, the dCas9-HDAC4 fusion protein was stabilized via removal of dTAG from the medium. After 2 more days, the cells were differentiated for 2 days via 2i/LIF-withdrawal and stained for Xist RNA using FlowFISH. Cells were sorted depending on Xist levels into Xist^Neg^ (bottom 15%) and Xist^High^ (top 15%) populations. Counts in the different populations were assessed via NGS. (b) Cell culture work flow for the TFiMini screen. Icons below the timeline indicate the point at which cells were harvested for the Selection and 2iSL populations. (c) Workflow of the design of the TFiMini library. The promoters outside of the *Xic*, included in the TFi library, were filtered based on their enrichment in the Xist^High^/Xist^Neg^ comparison (Wald.p≤0.2). Non-enriched published TF regulators were added, despite missing the significance threshold. The top 8 guides were selected by absolute fold change in the Xist^High^/Xist^Neg^ comparison in the TFi screen. Lastly, guides targeting Xist-controlling REs and the FIREWACh minimal promoter were added as controls^15,52^. (d) Xist RNA expression assessed by flow cytometry following FlowFISH staining. Additionally to the library (TFiMini) the plot also depicts an undifferentiated (2iSL) and a +dTAG control (CRISPR system not induced). (e) Expression of Xist and the naive pluripotency factors *Prdm14* and *Nanog* during the TFiMini screen assessed by RT-qPCR. The black bars denote the mean between two replicates (dots). (f) The sgRNA distribution width of the normalized counts in the different populations of the TFiMini screen, separated by targeting (left) and non-targeting guides (right). The score is given by the subtraction of the log_2_-transformed values for the 90^th^ and 10^th^ percentile. (g) Correlation between the two replicates in the different populations of the TFiMini screen (1,270 guides). The Pearson correlation coefficient is indicated in the plots.

**Extended Figure E4:**
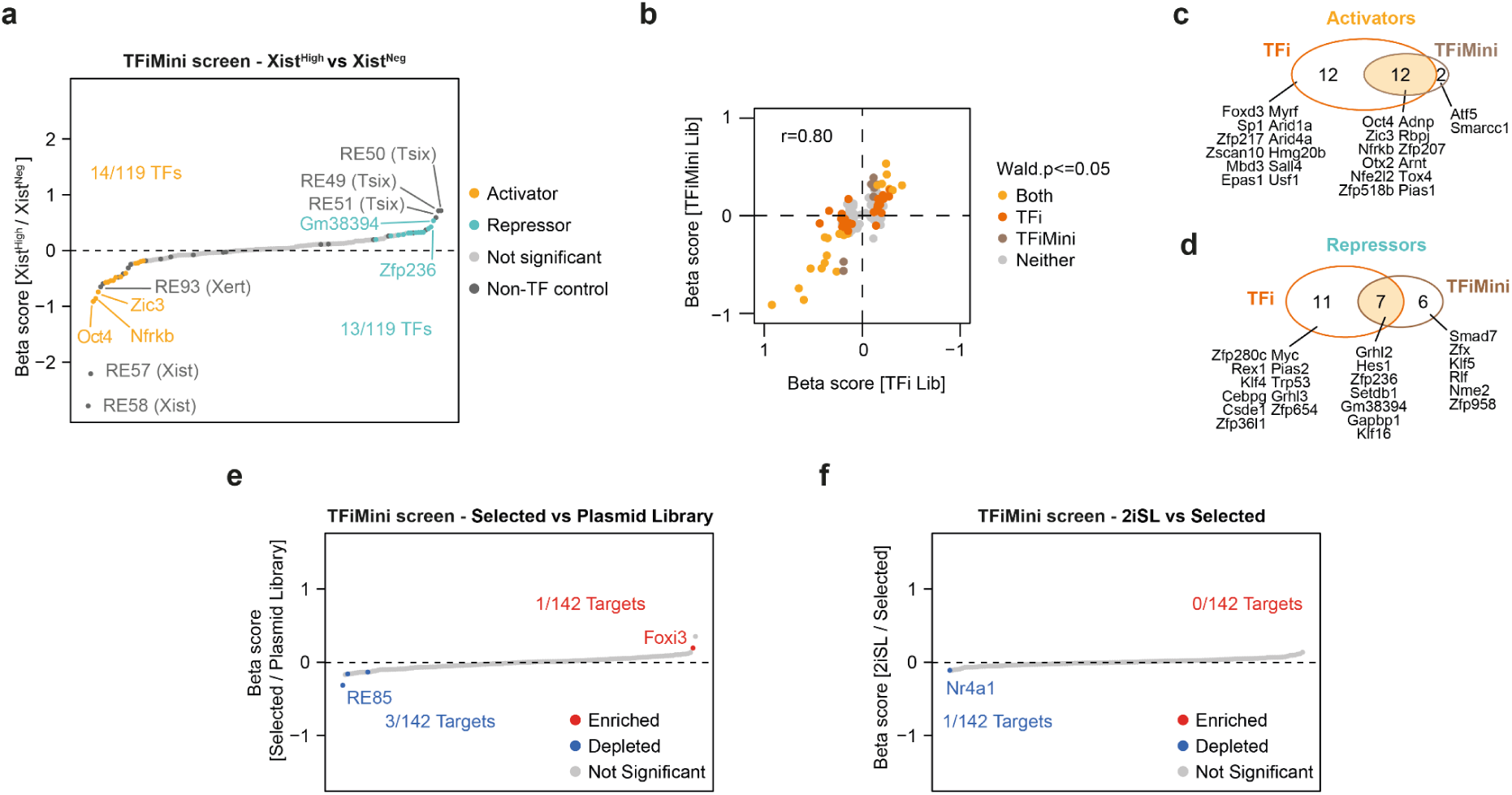
(a) Comparison of target abundance between Xist^High^ and Xist^Neg^ populations in the TFiMini screen. Significantly enriched or depleted targets are colored in teal or orange respectively (*MAGeCK mle*, Wald.p≤0.05). Published non-TF controls are colored in dark gray. (b) Correlation between the Xist^High^/Xist^Neg^ comparisons of the TFi and TFiMini screens (119 shared TF genes). Significantly enriched targets (Wald.p≤0.05) are colored in light orange (both screens), dark orange (TFi screen only) or brown (TFiMini screen only). The Pearson correlation coefficient is indicated in the plot. (c-d) Venn diagram showing the overlap in significant activators (c) and repressors (d) (Wald.p≤0.05) between the Xist^High^/Xist^Neg^ comparisons of the TFi and TFiMini screens. (e) Ranked plot depicting enriched and depleted targets between the Selected population and the plasmid library in the TFiMini screen. Significance was assessed using *MAGeCK mle* (Wald.p≤0.05). (f) Ranked plot depicting enriched and depleted targets between the 2iSL and Selected populations in the TFiMini screen. Significance was assessed using *MAGeCK mle* (Wald.p≤0.05).

**Extended Figure E5:**
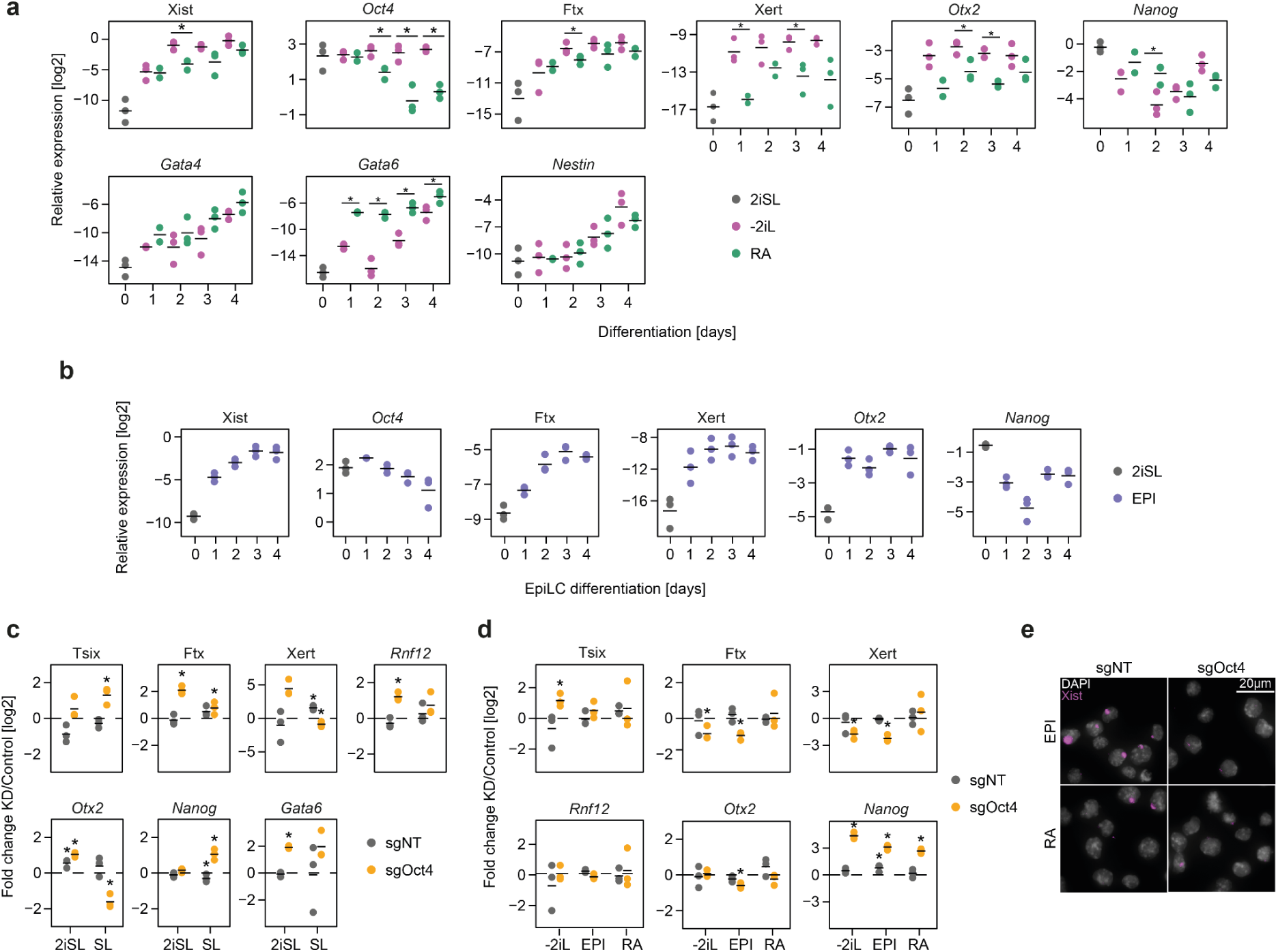
(a) RT-qPCR results for Xist, *Oct4* and other selected genes during 2iL-withdrawal only (-2iL) and in the presence of retinoic acid (RA). Significant difference between the differentiation protocols are marked with a black asterisk (*unpaired t-test*, p≤0.05). The black bars denote the means across three replicates. (b) RT-qPCR results for Xist, *Oct4* and other selected genes during differentiation toward EpiLCs. The black bars denote the means across three replicates. (c) Effect of *Oct4* knockdown on the expression of selected Xist regulators in 2iSL or SL conditions, assessed via RT-qPCR. The results are shown as a log_2_ fold change between the control and knock down conditions. Significant differences are marked with a black asterisk (*unpaired t-test*, p≤0.05). The black bars denote the means across three replicates. (d) As (c), but after two days of differentiation via 2iL-withdrawal (-2iL), towards EpiLCs (EPI) or in the presence of retinoic acid (RA). (e) Microscope images corresponding to EPI and RA samples from Fig. 2c.

**Extended Figure E6:**
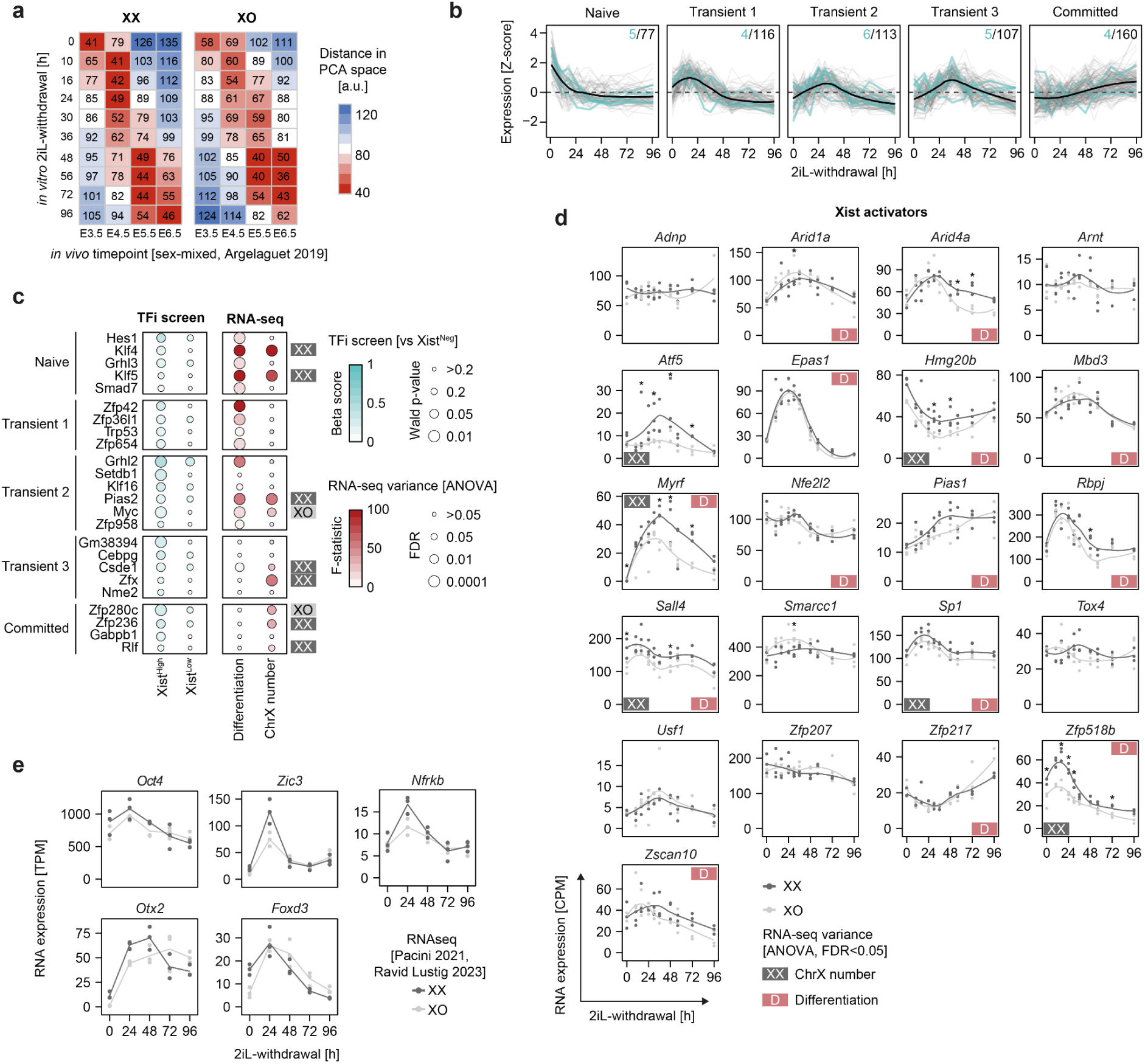
(a) Pairwise distance in PCA space between published scRNA-seq data of developing mouse embryos and the XX or XO 2iL-withdrawal RNA-seq time course^48^. Sex-mixed cells from the *in vivo* samples were filtered for ICM and epiblast cells and aggregated as pseudo-bulk. (b) Clustering of z-score transformed XX expression dynamics for all TF genes included in the TFi screen (groups based on XX and XO cells). Xist repressors are shown in teal. The total numbers of genes and repressors per cluster is indicated in the top right. (c) Dotplot depicting the TFi screen results versus the Xist^Neg^ population (left, *MAGeCK mle*) and dependence of the RNA-seq data on differentiation or X-chromosome number (right, *two-way ANOVA*) for the identified Xist repressors. The directionality for the X-chromosome effect is indicated next to the plot (FDR≤0.05). Expression clusters of the different activators are indicated on the left. (d) RNA expression dynamics of identified Xist activators. Difference between XX and XO cells was assessed per time point using *DESeq2* (Wald.FDR≤0.05)^51^. Results from the ANOVA analysis are indicated as D (affected by differentiation) or XX (affected by X-chromosome number). (e) RNA expression dynamics of selected Xist activators in a published 2iL-withdrawal time course^36,40^.

**Extended Figure E7:**
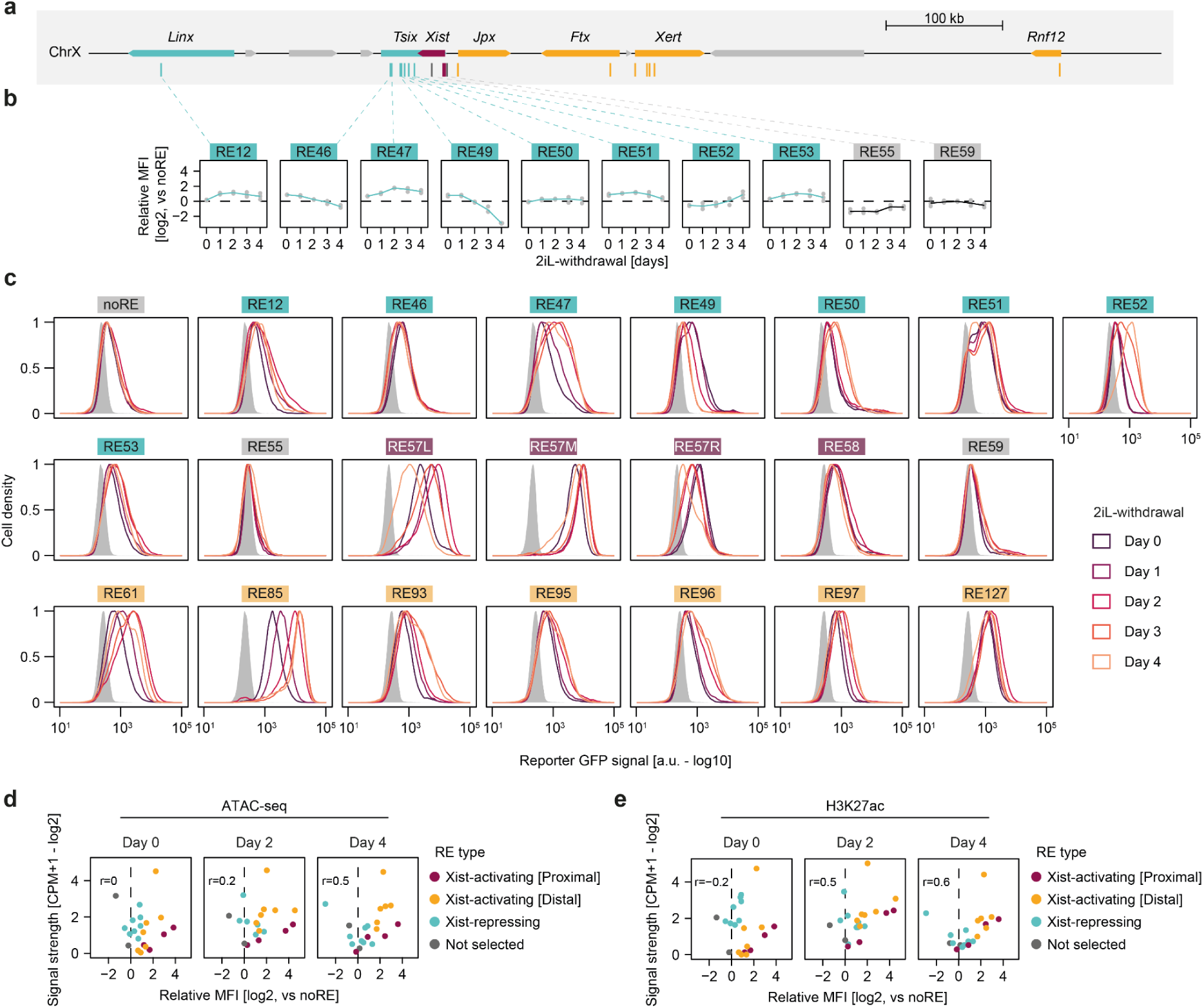
(a) Schematic outlining the genomic region surrounding the *Xist* locus. Xist regulators are shown in teal (repressive) or orange (activating). Xist-controlling REs included in the reporter time course are shown in teal (Xist-repressive), gray (not selected for reporter screens), burgundy (proximal) or orange (distal). (b) Reporter expression dynamics for Xist-repressive (teal) and non-selected REs (gray). Relative MFI was calculated as the log_2_ fold change between GFP levels in the indicated reporter and the noRE control. Mean (line) of n=3 biological replicates (dots) is shown. (c) Reporter GFP signal assayed by flow cytometry for the data summarized in (b) and Fig. 4c. One representative replicate is shown. The gray shade corresponds to non-fluorescent control cells. (d) Correlation between the reporter time course (mean of 3 biological replicates) and published chromatin accessibility data (mean of 2 biological replicates)^15^. The ATAC-seq data was generated in a female heterozygous TX Δ*Xic* deletion line. The assayed counts thus correspond to the Xi. The Pearson correlation coefficient is indicated in the plots. (e) As (d), but using published CUT&Tag data targeting H3K27ac^15^.

**Extended Figure E8:**
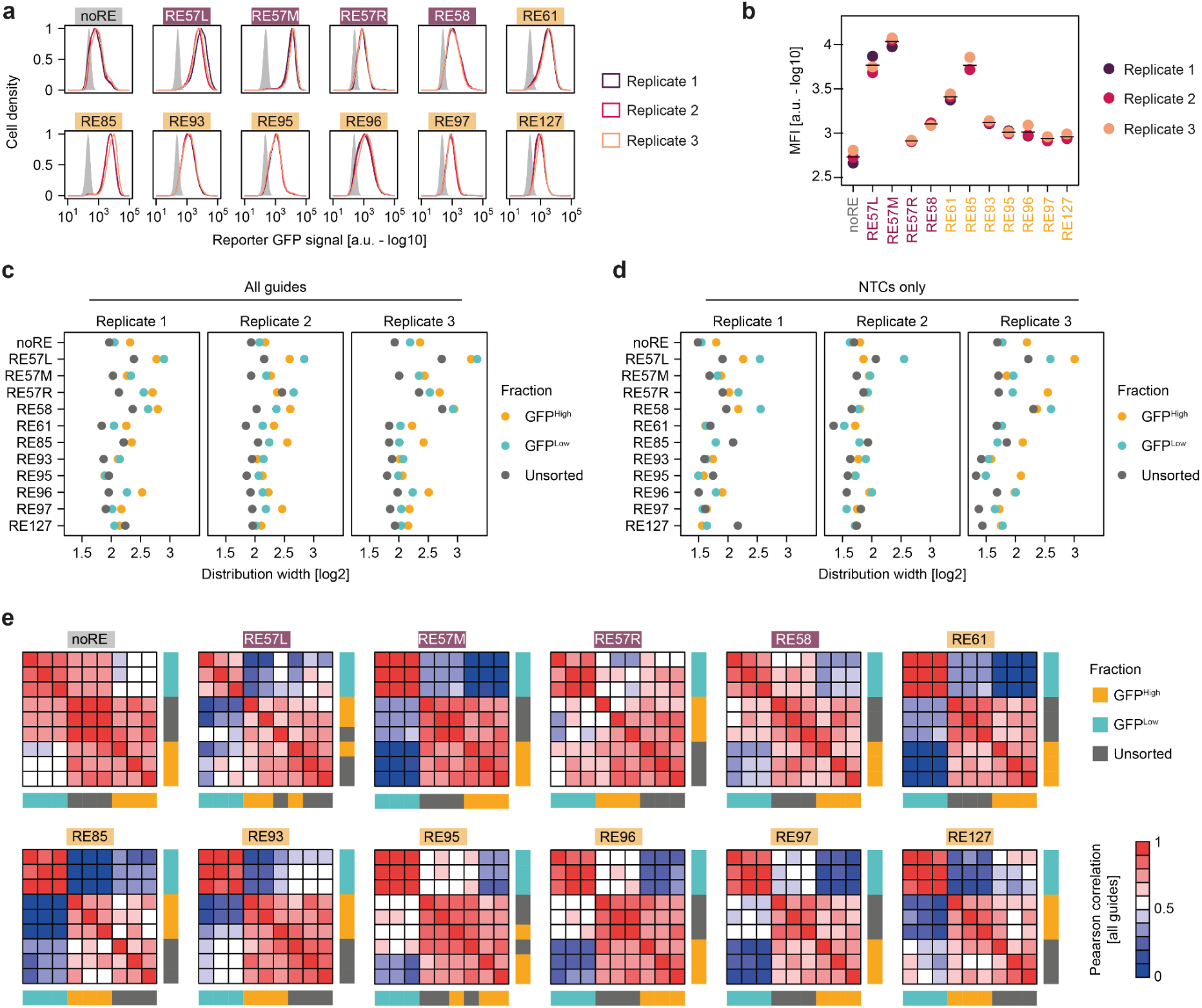
(a) Reporter GFP signal during the reporter screens. The gray shade corresponds to non-fluorescent control cells. (b) Quantification of the data depicted in (a) as the mean fluorescence intensity. The black lines denote the geometric mean across three replicates. (c) SgRNA distribution width of the normalized counts in the different populations of the reporter screens, including all targeting guides. The score is given by the subtraction of the log_2_-transformed values for the 90^th^ and 10^th^ percentiles. (d) As in (a), but including only non-targeting controls. (e) Correlation of the normalized counts between all replicate samples per reporter screen. The populations are indicated by colored boxes. The axes are ordered by hierarchical clustering.

**Extended Figure E9:**
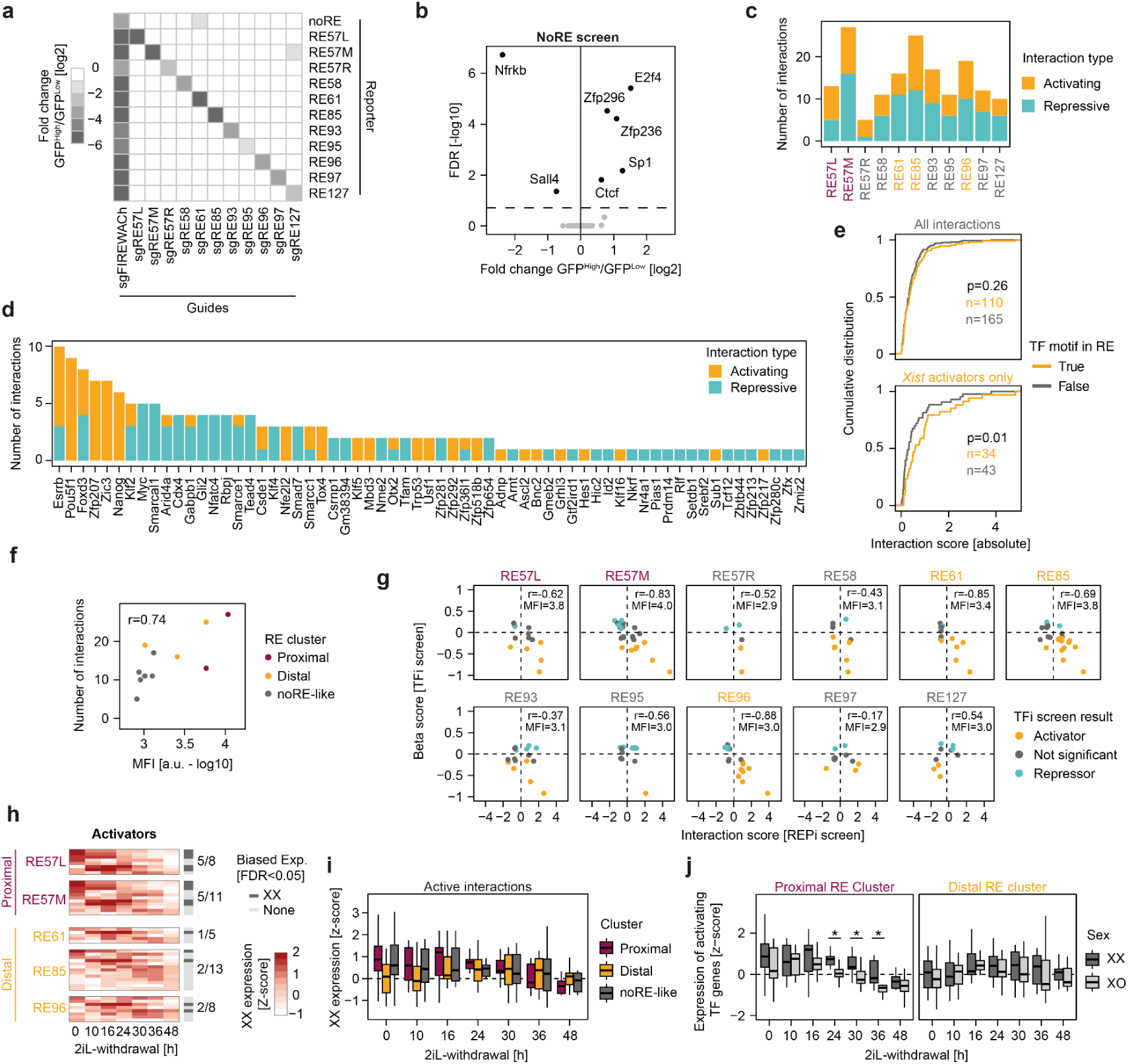
(a) Effects of positive controls in the reporter screens. Fold change of the normalized counts between the GFP^High^ and GFP^Low^ populations for guides targeting the FIREWACh minimal promoter or the respective RE sequences. The heatmap denotes the mean across 3 biological replicates. (b) Enrichment of TF targets in the noRE screen. All 7 significant hits (labeled) were removed from further analysis (*unpaired t-test*, FDR≤0.2). (c) Number of activating and repressive interactions per reporter line. (d) Number of activating and repressive interaction per TF target. (e) Cumulative distribution of the absolute interaction scores showing the correlation between the presence of a binding motif and interaction strength. The bottom panel shows only interactions of Xist activators from the TFi screen (Xist^High^/Xist^Neg^ comparison). Significance was calculated using a *rank-sum Wilcoxon test* and is indicated in the panels. The number of tested interactions with (orange) and without (gray) a motif is indicated in the panels. (f) Correlation between base signal strength of the reporter line and the number of identified interactions. The signal strength was quantified as the MFI during sorting. The Pearson correlation coefficient is indicated in the plot. The reporter screens are colored by RE cluster (see Fig. 4g). (g) Correlation between reporter screen hits and the TFi screen separated by reporter line (Xist^High^/Xist^Neg^). The Pearson correlation coefficient is indicated in the panels. The mean MFI during sorting is also indicated in the panels. (h) Z-score transformed XX expression dynamics of activators for selected reporter lines during the RNA-seq time course (Figure 3). TF activators are ordered via hierarchical clustering. Interaction partners with XX-biased expression are marked on the right (*ANOVA*, FDR≤0.05) (i) Z-score transformed XX expression dynamics of TF genes activating a reporter line located in either the proximal, distal or noRE-like PCA clusters (Fig. 4g). The central line represents the median and the edges of the box represent the range from the 25^th^ to the 75^th^ percentiles. (j) As in (e), but comparing XX and XO expression for TF genes activating a proximal (left) or distal (right) reporter line (for cluster see Fig. 4g). Significance is marked by a black asterisk (r*ank-sum Wilcoxon test*, p≤0.05).

**Extended Figure E10:**
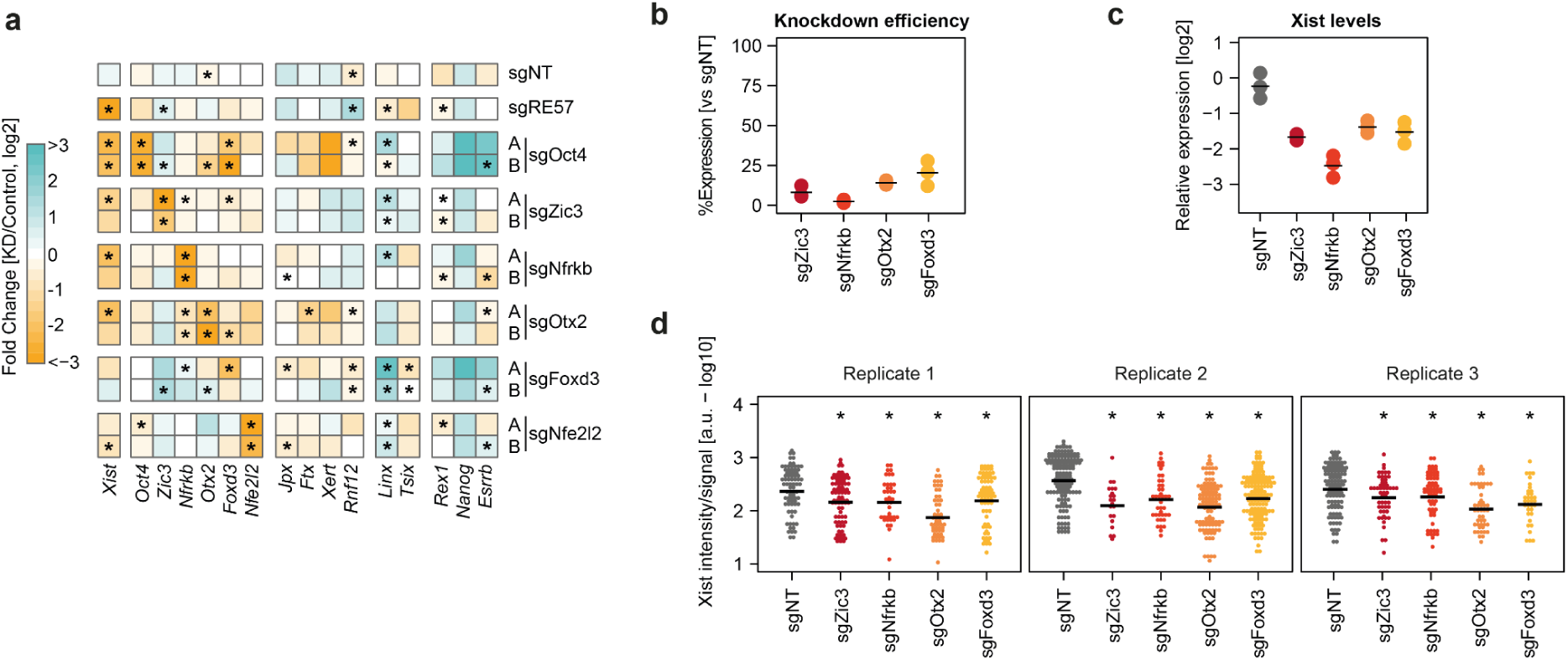
(a) Effect of knockdown of selected Xist activators via the HDAC4 CasTuner system on the expression of Xist and various Xist regulators assessed by RT-qPCR. Guides targeting RE57 were included as a positive control for Xist. The heatmap shows log_2_ fold changes of the relative expression between the KD and the +dTAG control samples (mean of 2 biological replicates). Two independent multiguide plasmids (A+B) were used per target. The black asterisk denotes significance (*unpaired t-test*, p≤0.05). (b) Knockdown efficiency during the RNA-FISH and CUT&Tag experiment assessed by RT-qPCR (see Fig. 5). The data is quantified per replicate compared to the non-targeting control. The black bar denotes the mean of three replicates. (c) Total Xist levels during the RNA-FISH and CUT&Tag experiments assessed by RT-qPCR (see Fig. 5). The black bar denotes the mean of three replicates. (d) Effect of TF gene knockdown on the RNA-FISH intensity per Xist signal, separated by biological replicate (see Fig. 5f). Xist signals were detected and quantified via an automated analysis pipeline. The black bar denotes the geometric mean per sample (21-150 Xist signals per sample). The black asterisk denotes significance compared to the sgNT control (*rank-sum Wilcoxon test*, p≤0.05).

**Extended Figure E11:**
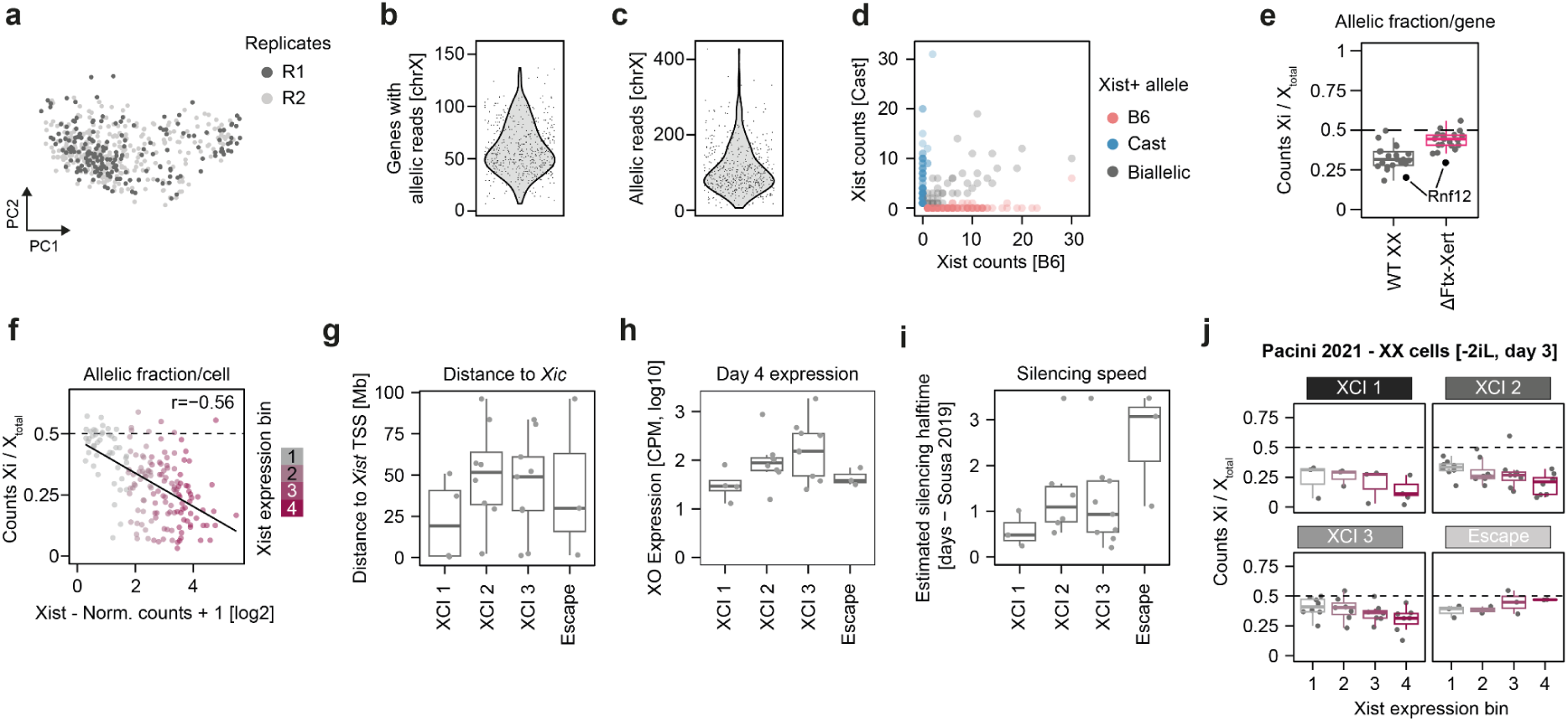
(a) PCA depicting single-cell transcriptomes included in the analysis separated by replicate (502 cells). (b) Number of X-chromosomal genes with allelic reads per cell (502 cells). (c) Total number of X-chromosomal allelic reads per cell (502 cells). (d) Raw number of allelic Xist counts for Xist+ cells separated by allele (252 cells). Biallelic cells (Xist allelic ratio 0.2-0.8) are shown in gray and were removed from further analysis (41 cells). (e) Allelic ratio of X-chromosomal genes, comparing wildtype and Δ*Ftx*-*Xert* cells. All genes included in Fig. 6h-i are shown (25 genes). The central line represents the median and the edges of the box represent the range from the 25^th^ to the 75^th^ percentiles. (f) Correlation between chrX allelic fraction and normalized Xist counts for wildtype Xist+ cells (178 cells). The Pearson correlation is indicated in the plot. A linear fit is depicted as a black line. The cells are colored by the Xist expression bin. (g) Genomic distance to the *Xist* TSS for X-linked genes separated by XCI group (see Fig. 6h-i). Gray dots denote individual genes. The central line represents the median and the edges of the box represent the range from the 25^th^ to the 75^th^ percentiles. (h) As in (g), but denoting expression of the genes in XO cells at day 4 of 2iL-withdrawal (data from Fig. 3), corresponding to the base expression in the absence of XCI. (i) As in (g), but showing estimated silencing half-lifes obtained in a previous study following induced XCI^54^. (j) Allelic expression for genes and XCI groups included in Fig. 6h-i for published scRNA-seq data (-2iL, day 3). Xist+ cells were binned according to normalized Xist expression (134 cells). Individual genes are shown as gray dots. The central line represents the median and the edges of the box represent the range from the 25^th^ to the 75^th^ percentiles.

## Supplementary tables

Supplementary Table S1. TFi/TFiMini CRISPR library design and screen analysis. Related to Figure 1 and Extended Figures E1-E4.

Supplementary Table S2. RNA-seq time course analysis. Related to Figure 3 and Extended Figure E6.

Supplementary Table S3. Reporter CRISPR screen analysis. Related to Figure 4 and Extended Figure E7-E9.

Supplementary Table S4. CUT&Tag and RNA-FISH in knockdown lines. Related to Figure 5 and Extended Figure E10.

Supplementary Table S5. Allelic scRNA-seq analysis. Related to Figure 6 and Extended Figure E11.

Supplementary Table S6. Experimental materials. Details utilized cell lines, primers, plasmids, antibodies and FISH probes.

